# Identification of low threshold off-target activation pathways during stimulation of carotid baroreceptor afferents in swine

**DOI:** 10.64898/2026.01.02.697436

**Authors:** Ashlesha Deshmukh, Kevin Cheng, Megan L. Settell, Bruce E. Knudsen, Maria LaLuzerne, Julie Savage, Kathryn Turk, Constantinos Tsiptsis, Kshirin Anand, Roger Schultz, Ricardo Sanchez, Annabelle Olson, Aaron Suminski, Andrew J. Shoffstall, Warren M. Grill, Nicole A. Pelot, Kip Ludwig

## Abstract

**Objective:** Electrical stimulation of the baroreceptors pathways at the carotid sinus bulb – known as baroreflex activation therapy (BAT) – is intended to change autonomic tone and ultimately reduce blood pressure and heart rate. BAT is pre-market approved by the United States Food and Drug Administration (FDA) for the treatment of heart failure and received an FDA humanitarian device exemption for drug resistant hypertension. However, responder rates are limited by side-effects including numbness in the head and neck, altered speech, respiratory constriction, dry cough, vomiting, and altered sensory and motor function of the tongue. We hypothesized that these side-effects are driven by activation of other nearby nerve fibers of similar or lower threshold than the carotid sinus nerve. In this study, we sought to identify the neural sources responsible for off-target muscle activation contributing to these side-effects. These sources would inform strategies mitigating off-target activation in BAT therapy.

**Methods:** Domestic swine were used in this work as the diameter and thickness of the swine carotid artery are closer to human than those of canine models. A monopolar disk electrode mimicking the clinical CVRx® Neo electrode was surgically placed proximal to the carotid bifurcation with the position optimized for stimulation dose responsive changes in blood pressure. Evoked responses were recorded during dose response testing from multiple neck muscles, and the corresponding off-target nerve pathways were identified by sequential transection of nearby nerves.

**Results:** The following activated off-target muscle groups and their corresponding nerve pathways were verified which included 1) the cricoarytenoid via recurrent laryngeal nerve, 2) cricothyroid via superior laryngeal nerve, 3) sternocleidomastoid via accessory nerve, and 4) the tongue via hypoglossal nerve. The constrictor muscle group was also activated through a more complex neural pathway.

**Conclusion:** We identified multiple sources of therapy-limiting side-effects of BAT in the swine model. These results will help guide the design of improved stimulation electrodes, surgical placement, and parameter programming to reduce BAT-associated side-effects while increasing on-target activation in patients.

## INTRODUCTION

Hypertension, defined as arterial systolic/diastolic blood pressure (BP) greater than (140/90 mmHg), occurs in 48.1% of the U.S. population, i.e., 20 million adults (CDC, 2024). Of these, 20% are further classified as having treatment-resistant hypertension (RHT). RHT is defined as 1) uncontrolled BP (≥ 130/80 mmHg) despite concurrent prescription of 3 to 4 antihypertensive drugs of different classes, or 2) controlled BP using more than four antihypertensive medications, with each drug administered at the highest tolerated dose (Carey et al., 2018). Approximately 23 million US adults have uncontrolled hypertension that is resistant to pharmacological solutions. Chronic hypertension can lead to heart failure (Messerli et al., 2017), with 6.7 million adults suffering from the disease and a 5-year survival rate of 43% after first hospitalization (Tribouilloy et al., 2008). This clinical need led to the development of novel device-based therapies such as Baroreflex Activation Therapy (BAT), which currently has premarket approval (PMA) from the U.S. Food and Drug Administration (FDA) and European CE mark approval to treat heart failure (Abraham et al., 2015a; Gold et al., 2016; Kumar et al., 2023; Zile et al., 2020) in patients who stopped responding to traditional drug therapies (Bakris et al., 2012; Halbach et al., 2020).

BAT is based on electrical activation of baroreceptors, which are mechanoreceptors that innervate the carotid artery bifurcation and aortic arch. Baroreceptors measure tonic and phasic changes in BP to maintain constant blood perfusion in the body. BP fluctuations are measured at the carotid sinus and transmitted via the carotid sinus nerve (CSN), a branch of the glossopharyngeal nerve (GPN), to the nucleus of the solitary tract (NTS) in the brainstem. Changes in the NTS signaling cause an increased parasympathetic outflow to the heart and a reduction in sympathetic tone, resulting in the dilation of blood vessels and a reduction in heart rate (HR). In combination, these changes can cause a sustained reduction in blood pressure (Abraham et al., 2015b; Victor, 2015). Preclinical results laid the foundation for clinical studies, which showed significant and sustained reductions in BP with additional studies exploring safety and efficacy in heart failure (NCT01471860, NCT01720160, NCT02627196).

These clinical studies (Bisognano et al., 2011; Scheffers et al., 2010; Victor, 2015) led to regulatory approval and early adoption of BAT in clinics. However, many patients also reported side-effects such as cough, throat pain, dysphonia, difficulty swallowing, gagging, and difficulty breathing that were intolerable for extended time periods, severely limiting the therapy efficacy (Jordan et al., 2024). One such clinical study, using CVRx’s FDA-approved electrode ‘Neo’, showed a - 16.9 ± 15 (avg. ± SD) mmHg decrease in systolic BP in 18 patients. However, when stimulation therapy dosing amplitude was turned down to tolerable levels to avoid side-effects, BP decreased by only -6.3 ± 7 mmHg (Heusser et al., 2016).

To minimize side-effects of BAT with revised electrode design or placement, it is important to understand the neuroanatomical pathways responsible for off-target effects relative to the targeted carotid sinus bulb. Importantly, motor nerves are frequently activated at currents lower than those required to activate smaller diameter fibers, and have been implicated as the source of limiting side effects for BAT and vagus nerve stimulation (Blanz et al., 2022; Nicolai et al., 2020; Tosato et al., 2006, 2007; Yoo et al., 2013). The swine model was chosen for this study due to its similarity to human anatomical dimensions, as distances are key for electrical activation (Pelot et al., 2020; Settell et al., 2020). We first used angiography to visualize the arterial structures and microdissection to identify likely neuroanatomical structures driving off-target activation in the vicinity of the carotid bifurcation in swine. Next, dose-response curves were generated while applying BAT through an electrode intended to mimic the clinical Neo up to amplitudes sufficient to generate reliable changes in HR and BP to understand the therapeutic window between effective BAT and activation of off-target nerves. Then functional confirmation of the relevant off-target anatomy was performed using sequential transections of each putative nerve pathway and electromyography from its terminal muscle grouping. By identifying the neuroanatomical substrates responsible for off-target activation during BAT and assessing their dose-response curves with respect to clinical BAT, these results will inform future surgical placement strategies and electrode designs/stimulation patterning to mitigate the impact of side-effects on this promising therapy.

## METHODS

### Swine

The swine model was used in this study to understand the on-vs. off-target pathways of BAT because the pig anatomy is similar to humans in terms of carotid artery size and thickness, nerve sizes, anatomical landmarks, and complexity of fascicular organization of nerves (Mathern et al., 2022; Settell et al., 2020). A total of fifteen swine were used for this study. Thirteen were used in functional studies (8 M/5 F; 37.38 ± 4.00 kg, mean ± SD). Of these thirteen, five were used in pilot experiments for refining surgical technique, optimizing placement for reliable cardiac responses during BAT, visual identification of activated off-target muscles, and microdissections to identify relevant nerve pathways. These five swine are not included in dose-response curve analyses; data from Swine 6-13 were used to analyze dose-response curves for the on- vs. off-target framework (swine-specific information in Supplemental table 1). Two swine were used for in vivo angiographic imaging and postmortem micro-computed tomography (microCT), bringing the total number of swine to fifteen. All swine were housed individually (21°C and 45% humidity) with ad libitum access to water, and were fed twice daily until 12 hours prior to the surgical procedure. All study procedures were approved by the University of Wisconsin-Madison Institutional Animal Care and Use Committee (IACUC) and were conducted under the guidelines of the American Association for Laboratory Animal Science (AALAS) in accordance with the National Institutes of Health (NIH) Guidelines for Swine Research (Guide for the Care and Use of Laboratory Animals).

### Imaging

Angiography was performed in n=2 swine to create a 3-D reference map of the carotid bifurcation, carotid sinus bulb, and the connected vasculature for illustrative purposes. These data were also used to supplement subsequent microCT images of “en bloc” carotid samples to understand the distances to on- and off-target nerve anatomy.

#### Angiography

All angiographic image acquisitions were conducted under anesthesia (1-3% inhaled isoflurane in conjunction with 2-100 mcg/hr constant infusion of fentanyl, dosed to effect). The swine were placed in a dorsal recumbent position with the head and neck under the angiographic imaging arm. The endovascular structure of the carotid arteries was visualized using computed tomographic angiography (CTA). Intra-arterial iodinated contrast [OMNIPAQUE (iohexol) Injection 300 mgI/mL] was loaded in a syringe for introduction. The common carotid artery was exposed using surgical cutdown for access near the sternum. The artery was cannulated using a MEDLINE 7 French (Fr) catheter. The contrast was manually introduced through the cannula using a 60 cc/ml Terumo syringe and a 3D volume scan was acquired using established protocols (Edwards et al., 2022). The angiography files were analyzed in 3D Slicer 5.6.1 by loading the 3D volume DICOM files and using the segmentation and volume rendering modules for visualization.

#### MicroCT

Postmortem dissections were performed to isolate the carotid bifurcations and the surrounding off-target nerves in an “en bloc” technique. The common carotid artery, with the vagus nerve and the sympathetic trunk attached, was isolated ∼2-4 cm caudal to the bifurcation and traced cranially. The off-target nerves (superior laryngeal nerve, accessory nerve, hypoglossal nerve, vagus nerve) and the glossopharyngeal nerve were traced from their terminal muscles back to the carotid bifurcation and labeled using color coded dye. The carotid bifurcation and the surrounding connective tissue (∼1 cm) were kept unperturbed to maintain the small neural branches and orientation of the off-target nerves closest to the stimulation location. The dissected nerves and arteries were cut at their distal ends, and the entire structure was transferred to a gridded acrylic board and glued down while aiming to maintain its in vivo orientation. This “en bloc” sample was then immediately transferred to 10 % neutral buffered formalin for immersion fixation. Formalin-fixed samples from UW Madison were shipped to Case Western Reserve University in 1x phosphate buffered solution (PBS). These samples were then stained in phosphotungstic acid (PTA) and scanned using Scanco® µCT100 instrument as per previously published protocols (R. Upadhye et al., 2025; A. Upadhye et al., 2025; A. R. Upadhye et al., 2025). Scanned image volumes were visualized using 3D Slicer and manual segmentation was carried out using the Slicermorph extension. Desired structures were then highlighted using the Volume Rendering module.

### Surgical

#### Anesthesia

For induction, each animal was given an intramuscular injection of Telazol (6 mg/kg) and xylazine (2 mg/kg). Animals were then ventilated and maintained under 1-3% inhaled isoflurane and 2-100 mcg/kg/hr of 10% fentanyl in lactated Ringer’s solution for the remainder of the experiment. To minimize the effect of isoflurane on autonomic feedback circuits for blood pressure control, the inhaled anesthetics were reduced to 1.5-2% isoflurane in O_2_ during functional data collection. Anesthetic depth was verified and maintained throughout, determined via vitals monitoring, jaw tone, and blink reflex. All vital signs—including temperature, HR, CO₂, and SPO₂—were continuously collected and recorded every 10 minutes.

### Surgical cutdown

The surgical approach to identify the carotid artery branches of interest for BAT and possible off-target nerve pathways was refined in a pilot cohort of 5 animals during in-vivo and post-mortem dissection studies. The surgical protocol used for BAT functional studies is reported below.

#### Carotid artery cutdown

The swine was placed on its back. A 15-20 cm incision was made approximately 2 cm lateral to midline along the lateral aspect of the sternohyoid muscle on the swine’s right side. Underlying tissues (e.g., fat, connective tissue, parotid gland, mandibular gland) were bluntly dissected dorsally along the lateral margin of the trachea to the level of the esophagus. The cutaneous colli and omohyoid muscles were transected during the dissection so that the surgical pocket could be retracted laterally to maximize access to the carotid sheath and carotid bifurcation.

The carotid artery was mobilized from the level of the thyroid gland to approximately 2-3 cm cranial to the common carotid artery (CCA) bifurcation. Dissection cranial to the bifurcation was done with care to avoid damage to the plexus of small nerves while exposing the following off-target nerves.

#### Off-target nerves

The off-target nerves were identified as likely candidates driving muscle activation based on their proximity to the stimulation location and microdissection in pilot swine (Supplemental Figure 2,3 and 4). In functional studies, the well-coalesced trunks of these nerves were isolated closest to the stimulation location and transected to confirm as off-target pathways. The surgical protocol with relevant landmarks to identify these nerves is detailed below.

***Accessory nerve***: The external branch of the accessory nerve passes dorsal to the vagus nerve at the level of the nodose ganglion, courses laterally towards the sternocephalicus and brachiocephalicus muscles, and then divides into ventral and dorsal branches. The ventral branch innervates the sternocephalicus muscle and the dorsal branch innervates the trapezius muscles.
***Hypoglossal nerve:*** The hypoglossal nerve passes ventro-lateral to the glossopharyngeal and vagus nerves. It curves cranial-ventrally and inserts towards the base of the tongue.
***Superior laryngeal nerve (SLN):*** The SLN originates from the nodose ganglion and ascends medio-ventrally, either as a single nerve that divides into the external and internal branches or as two separate branches (Settell et al., 2020). The internal branch continues medio-ventrally and inserts into the medial lateral wall of the thyrohyoid membrane and innervates the upper larynx and constrictor muscles. The external branch ascends caudal-ventrally along the lateral aspect of the thyroid cartilage to innervate the cricothyroid muscles.
***Recurrent laryngeal nerve (RLN):*** The RLN courses along the medial lateral aspect of the tracheal cartilage in close proximity to the vagus nerve. It inserts into the larynx just posterior to the cricothyroid, coursing along the medial wall of the thyroid cartilage to innervate the cricoarytenoid muscles.

#### Muscles

Muscle groups activated during BAT were instrumented with bipolar EMG electrodes to record evoked muscle responses. The identified muscle groups which were surgically accessible were: 1) cricothyroid (CT), 2) cricoarytenoid (CA), 3) the inferior constrictors (CS), 4) tongue (TN, rostral), and 5) sternocephalicus (SC). The SC muscle in swine is equivalent to the sternocleidomastoid in humans and the thyropharyngeal muscle in swine is equivalent to the inferior constrictor muscle in humans (German et al., 2009; Getty, 1975; Sack et al., 1982). In the rest of this paper, the muscles will be referenced as sternocephalicus (SC) and constrictors (CS), respectively.

### Instrumentation and experimental design for functional studies

After completing the surgical cutdown of the arteries, nerves, and muscles described above, in vivo functional studies were conducted to record stimulation-driven decreases in blood pressure and EMG from activation of activated neck muscles.

#### Physiological recordings

Blood pressure (BP) and heart rate (HR) were collected using PowerLab systems (model 8/35, ADInstruments, New Zealand) and LabChart 8 software (ADInstruments, Sydney, Australia). All channels were acquired with a sampling rate of 1k samples/s. Physiological signals were aligned with electrical stimulation using a TTL-pulse synchronization signal between the stimulation and ADInstruments systems.

#### Blood pressure

A Millar catheter (Millar Inc., Houston, TX, model: MPR-500, size: 5F) was calibrated using a sphygmomanometer. Before calibration, the catheter was soaked in saline for 30 min, and the pressure gauge was adjusted to 0 for baseline reading. For calibration, ADinstruments Bridge Amp settings were adjusted for the BP acquisition channel in LabChart 8, wherein the voltage range was set to 20 mV to maximize resolution for small BP changes. The catheter was calibrated using the two-point reference technique using physiologically relevant points. It was then introduced through the femoral artery to the abdominal aorta (Figure 1A). Mean arterial pressure (MAP) was calculated and visualized in real-time using LabChart’s built-in measurement functions (1/3 maximum + 2/3 minimum of BP values per manufacturer’s instructions). The threshold for BP cycle (systole peaks) detection was 2 standard deviations from baseline (Supplemental material Figure 1). The calculated MAP was used for all analyses of BP.

**Figure 1:**
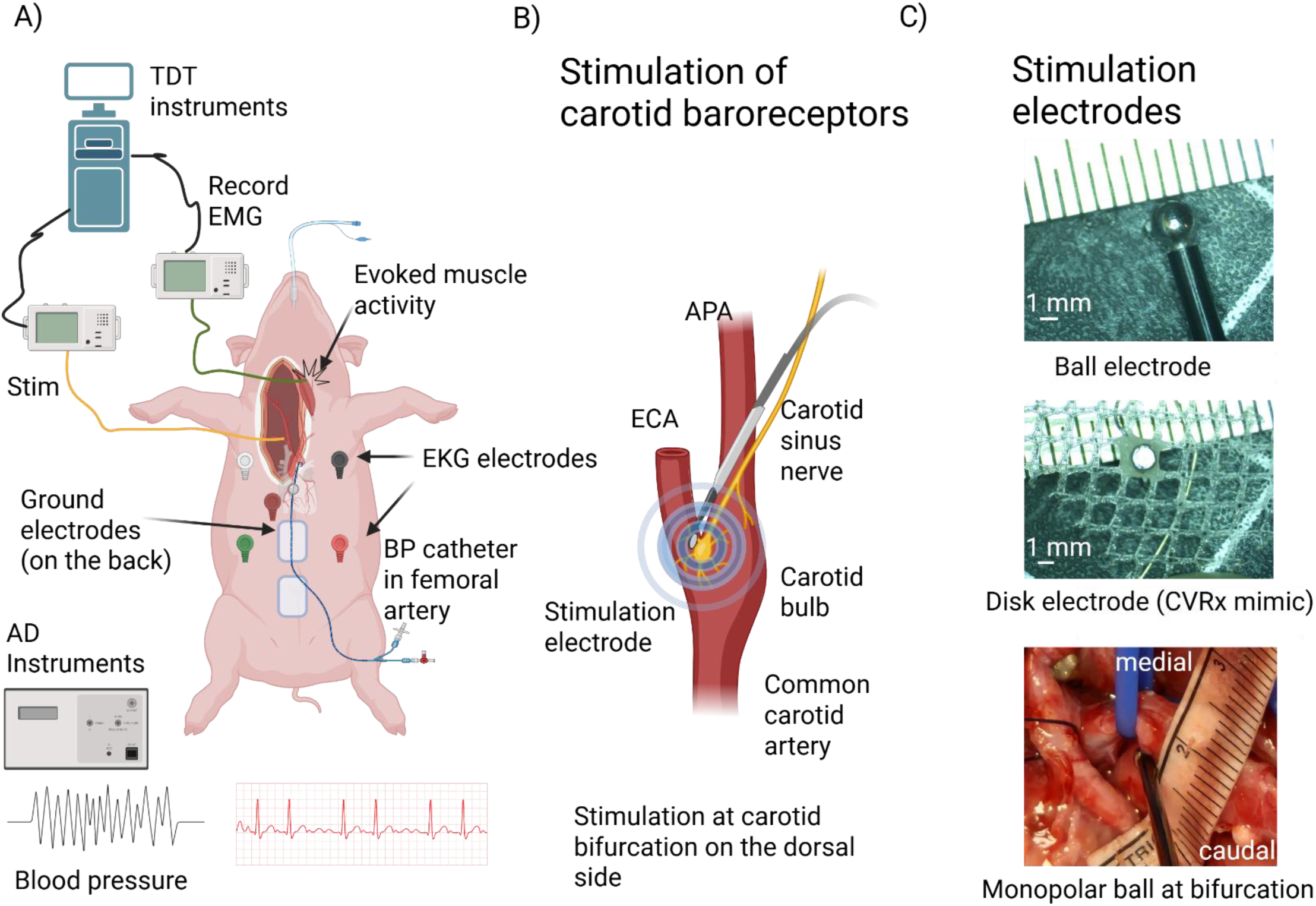
Experimental design. A) Schematic overview of surgical location and electrophysiological recordings. B) Location of stimulation electrode to target carotid baroreceptors. C) Stimulation electrodes: (top) ball electrode for mapping, (middle) 1 mm PtIr CVRx Neo mimic used to build dose response curves, (bottom) location of ball electrode placement in vivo during mapping of blood pressure responses to stimulation.

#### Heart rate

Electrocardiograms (EKG) were recorded using a traditional 5 electrode set-up (Figure 1A) using ADInstruments Dual Bio Amp with a digital bandpass filter (35-200 Hz). HR was calculated by detecting the QRS complex peaks using standard deviation thresholding and smoothed using a median filter with window width of 3001 samples or a 3 sec window.

#### Electromyograms instrumentation and recording

Electromyograms (EMG) recordings were conducted using Tucker Davis Technology (TDT) electrophysiology system (RZV10, TDT, Alachua Florida, USA). The evoked EMG was recorded from the CT, CA, SM, TN and the CS in a differential bipolar setup at a sampling rate of 25k samples/s. EMGs were recorded using paired fine stainless-steel wires (0.002” diameter) with approximately 2 mm of exposed tip (Rhythmlink, Columbia, SC, Product number: 221-28SS-550).

#### Stimulation instrumentation and recording

BAT stimulation was delivered using Tucker Davis Technology (TDT) electrophysiology system (IZV10, TDT, Alachua Florida, USA). Monopolar electrical stimulation was delivered with a stainless-steel grounding pad placed under the swine acting as the return electrode (Figure 1 A). The stimulation parameters tested were symmetric biphasic pulses with pulse widths (PW) (175, 250, 350 µs) at 25 Hz (Supplemental Table 1). These stimulation parameters were chosen to mimic the clinical BAT parameter range. Stimulation amplitude was titrated to generate reliable repeatable BP reduction. Every stimulation parameter was delivered for 30 secs, followed by two and half minutes or a longer recovery time for the baseline to return to pre-stimulation levels. See Supplemental Table 1 for swine-specific information. Stimulation mapping to find the location for BP decreases at the carotid bifurcation was carried out using a clinical ball electrode with a diameter of 3 mm and shaft length of 13 cm (MPM, Medical Supply, Freehold, New Jersey. Product number: LE-13-003) (Figure 1C, top and bottom). The on-target stimulation was delivered using an in-house manufactured 1 mm diameter Platinum (Pt) disk attached to a suture mesh for securing within tissue and backed with epoxy for insulating the back surface (Figure 1C, middle) intended to mimic the clinical Neo electrode.

### Data analysis and statistics

#### Evoked muscle activity

The evoked EMG was quantified using custom software (pyeCAP: https://github.com/ludwig822 lab/pyeCAP). The EMG responses were stimulus-triggered averaged over stimulation periods of 30 s and quantified as the root mean squared of the area under curve (AUC). The AUC integration windows were manually defined per muscle per swine to omit the stimulation artifact. All AUC values were normalized (per swine per muscle group) to compare responses across swine. The normalization was implemented using the min/max method where the maximum response was equated to 1. The normalized AUC values were plotted against stimulation amplitudes to generate dose-response curves. Lastly, evoked EMG signals at an amplitude above threshold were compared pre- and post-nerve transections to verify off-target nerve pathways.

#### Blood pressure and heart rate

Physiological data were exported using LabChart 8 in MATLAB file format and imported into pyeCAP (https://github.com/ludwig-lab/pyeCAP) for analysis. Stimulation pulses were synchronized with physiological responses to determine dose-dependent BP and HR effects. Baseline BP and HR were calculated as the average over 30 s before stimulation. The changes in BP and HR were calculated as the maximum change from baseline until 30 s post-stimulation to account for delayed BP changes beyond the stimulation period and normalized against baseline (max delta change value – baseline value, supplemental Figure 1) to account for swine-to-swine differences and any baseline drift over the course of the experiment (Figure 3A and B). Maximum BP and HR changes per animal were normalized to a 0 to 1 scale to facilitate cohort comparisons. An increase in BP from baseline or the minimum decrease from baseline was rescaled to 0, with 1 representing the maximum BP/HR decrease. Since this paper focuses on comparison of target effect (reduction in BP) and side-effect (activation of muscles), increases in BP were denoted as no effect or zero. Swine 8 was excluded from BP quantification and analysis due to lack of metadata about physiological recordings.

Statistical analyses were conducted using Python version 7 and SPSS version 28.0.1.0. Additional statistic data comparisons were conducted using scipy’s statsmodel library and statsmodel Api. Data visualization and analyses were done in Python using pandas, matplotlib, scipy, numpy, plotly, and seaborn libraries. To understand the extent of off-target muscle activation as a function of on-target responses, the normalized evoked muscle activity was compared against the normalized BP reductions. BP was treated as the independent variable (x) and EMG as dependent variable (y).

## RESULTS

The goal of this study was to understand the neuroanatomical sources of off-target muscle activation during therapeutic BAT. Towards that end, in the following results section we first outline the anatomical data proximal to the carotid bifurcation including angiography and post-mortem dissection to orient the reader. We then present dose response curves for HR and BP responses to verify optimal initial placement of the Neo mimic electrode for on-target responses prior to assessing off-target activation. Next, electromyography (EMG) is used to confirm concurrent activation of specific muscle groups implicated in reported off-target effects as a function of dose. Finally, the nerve pathways proximal to the bifurcation, that are the suspected sources of evoked neural activity ultimately resulting in the recorded muscle response are verified using transections.

### Anatomical documentation of swine carotid trifurcation and location for BP decrease

We performed microdissections (n=10) and angiography (n=2) to document the anatomical features of the right cervical common carotid bifurcation in swine. While microdissections informed the surgical approach for functional data, angiography enabled visualization of arterial networks that were difficult to trace without an extensive surgical cutdown which may disrupt relative positioning (Figure 2). The common carotid artery (CCA) was traced from the sternum (caudal) to the mandibular angle (cranial) to identify the carotid bifurcation of interest; in humans, the carotid sinus is located at the bifurcation between the internal carotid artery (ICA) and external carotid artery (ECA). The first small artery off the CCA was the superior laryngeal artery (SLA), which branched medial and travelled towards the esophagus (Figure 2A and E). The CCA split into a bifurcation cranial to the SLA, consisting of the ECA branching ventral-lateral from the CCA and the ascending pharyngeal artery (APA) branching dorsal-medial from the CCA. Compared to humans, in swine the APA serves as the main branching arterial structure replacing the ICA. However, the swine APA is slightly smaller in diameter ( ∼3 mm as compared to ∼6 mm) (Edwards et al., 2022). The ECA divided again cranially into a trifurcation of the lingual artery (LA), the facial artery (FA), and the mandibular artery (MA) (Duisit et al., 2017); this trifurcation was not traced in most specimens as it was outside the surgical window of interest. A small artery, the occipital artery (OA), branches off the APA near the CCA-APA-ECA bifurcation (Arikan et al., 2017; Kim et al., 2023).

**Figure 2:**
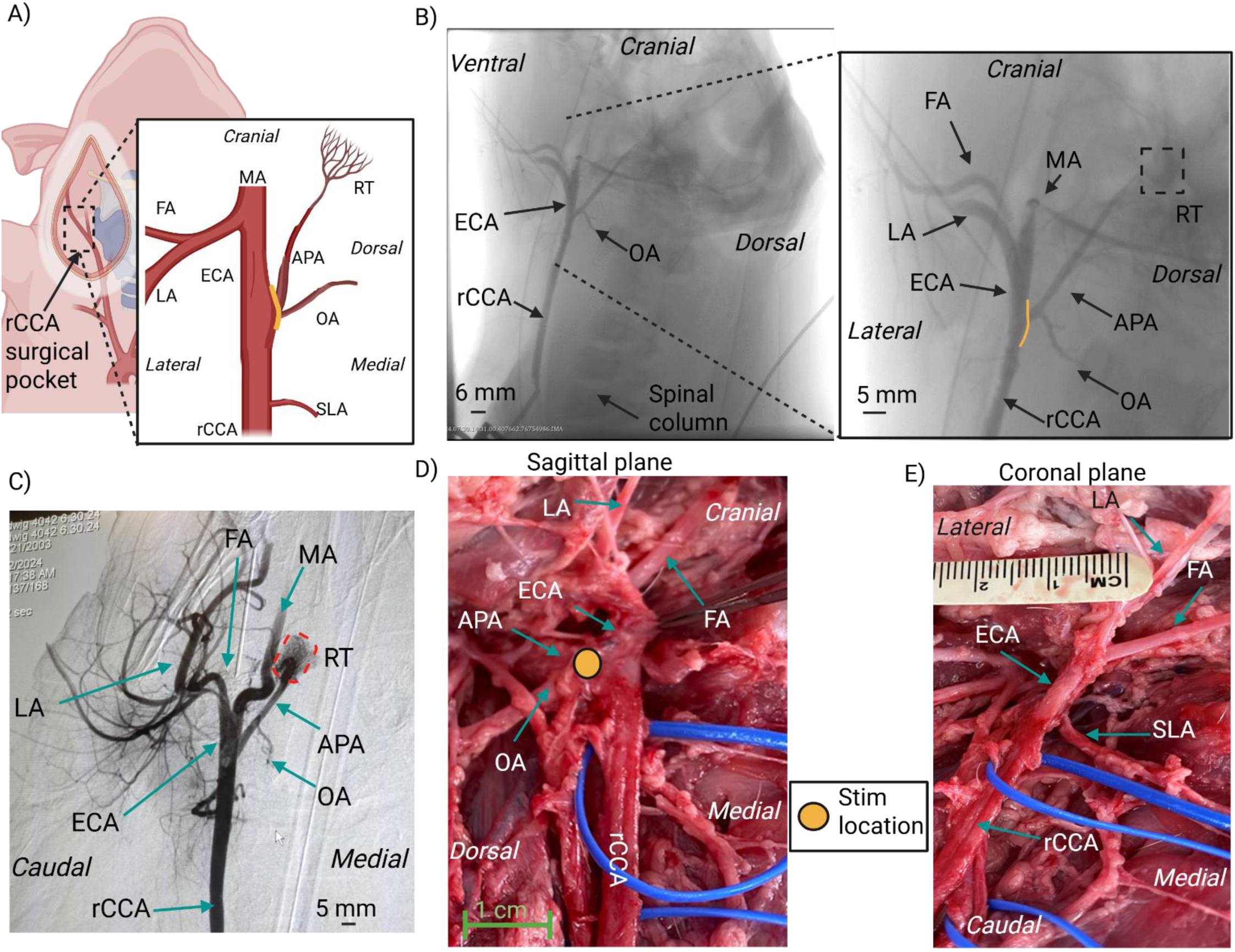
Anatomy of the swine carotid bifurcation. – A) Diagram of swine carotid anatomy where the CCA bifurcates into the ECA and the APA. SLA is the smallest in size and most rostral arterial branch that supplies the neck muscles on the medial side. The carotid bifurcation consists of the ECA that branches off laterally and the APA that branches off medially and courses dorsal and cranial. The APA gives off another smaller artery branch (OA) immediately, which runs medial and caudal. The ECA divides into three main branches: the FA, the LA, and the MA. The yellow line indicates the approximate stimulation location. B) Intra-operative visualization of angioCT data of right carotid artery with the contrast introducer needle at the caudal end of the CCA. Note that the right common carotid artery (rCCA) bifurcation pictured with the rCCA-ECA-APA branches was left unperturbed. The inset zoomed picture shows the APA coursing dorsal and cranial with the OA branching off near the bifurcation. The yellow line at the APA branching point shows the approximate identified locations for stimulation driven decrease in BP. C) Higher contrast intraoperative angioCT visualization of rCCA from a ventral view (coronal plane) showing all major arteries, i.e., the CCA, ECA, LA, FA, MA, OA, and importantly, the APA terminating in a vascular mesh identified as the RT. Other micro-arterial structures visualized in the surgical pocket at the CCA-ECA-APA bifurcation highlight the vascular complexity. D) Photo of a post-mortem microdissection (sagittal/right view) with the ECA retracted medially to visualize the APA diving dorsal. The OA branches off the APA near the CCA-ECA-APA bifurcation. The yellow circle on the APA at the dorsal side of the APA/ECA notch shows the stimulation location where BP reductions were recorded. E) Photo of a post-mortem microdissection (coronal/ventral view) of the carotid bifurcation showing the SLA branching off from the CCA and traveling medially and the ECA bifurcation point (split into FA and LA) being cranial to the SLA branching point. The APA is medial/dorsal and hard to visualize from this ventral view. APA – ascending pharyngeal artery, ECA – external carotid artery, FA – facial artery, LA – lingual artery, MA – maxillary artery, OA – occipital artery, rCCA – right common carotid artery, RT – rete mirabile, SLA – superior laryngeal artery.

The APA immediately traveled dorsally toward the foramen and was deemed impractical to trace further in microdissections. The APA was identified in the angiography scans by its branching location from the CCA and its distinguishable termination into a tiny mesh or plexus of arterio-arterial connections called the rete mirabile (RT) (Arakawa et al., 2007; Massoud et al., 1994). The RT was dorsal and cranial to the bifurcation (Figure 2B, C, D) at the base of the skull.

### Stimulation at on-target location evoked decreases in BP that depended on stimulation amplitude

To identify stimulation locations that evoked decreases in BP, we stimulated ∼± 1 cm caudal to cranial along the carotid bifurcation (CCA-ECA-APA) by moving the ball electrode in ∼ 2 mm increments while stimulating continuously at or above 4 mA. The location along the bifurcation that resulted in maximum decrease in mean arterial pressure (MAP) was identified in all pigs as the dorsal side of the bifurcation notch toward the trunk of the APA branching from the CCA (solid yellow line dorsal location in Figure 2B and yellow circle in Figure 2D). We designated this location as the on-target location for the remainder of the experiments and analyses.

Following the identification of the on-target location, a 1 mm disk electrode was sutured into place by placing a stitch using the electrode suture mesh (Figure 1C). This minimized electrode migration and maintained the distance between the on-versus off-target neural substrates for the remainder of the experiment; the electrode backing also provided insulation, as is used clinically to help direct the current toward the artery. Monopolar biphasic stimulation pulses mimicking clinical stimulation waveforms were delivered with varying stimulation amplitudes (1-10 mA) and fixed pulse widths to record stimulation dependent physiological responses. Stimulation produced obvious, repeatable dose-dependent decrease in mean arterial pressure (MAP, Figure 3). Muscles activated during stimulation that produced reductions in MAP (i.e., “therapeutic” stimulation) were visually identified in the surgical pocket and used to determine candidate nerve pathways for subsequent microdissections and the EMG placement in later functional experiments. MAP changes depended on stimulation amplitude in all swine while changes in the HR were more variable (Figure 3B and C, Supplemental Section Cohort blood pressure drops).

**Figure 3:**
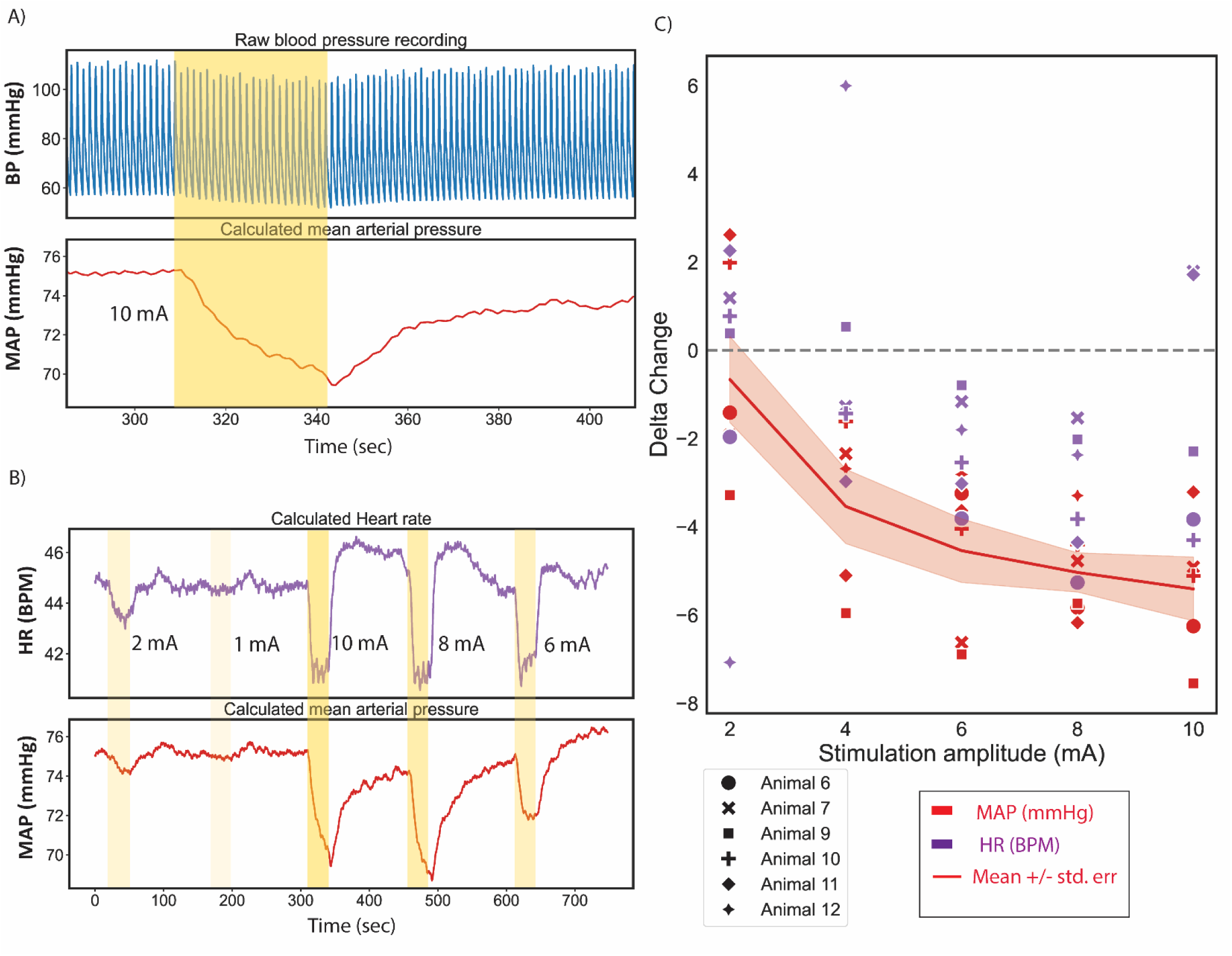
Stimulation at on-target location causes a decrease in blood pressure. A) Stimulation (10 mA, 150 μs PW, 25 Hz; ON during yellow shaded period) delivered through 1 mm disk electrode at carotid bifurcation caused a decrease in blood pressure (BP) (blue trace). Mean arterial pressure (MAP, red) calculated from the recorded BP, shows a decrease of ∼6 mmHg from the start of stimulation with a slow recovery/return to baseline. Data from Animal 6. B) Higher stimulation amplitudes (highlighted yellow bands with opacity representing stimulation intensity) caused a larger decrease in MAP (red) and heart rate (HR, in beats per minute (BPM); purple). Data from Animal 6. C) Cohort data for dose-dependent changes in MAP (red) and HR (purple). Changes were quantified as maximum change from baseline.

### Identification of possible off-target nerve pathways using microdissections

Microdissections in pilot swine were conducted to identify possible off-target nerve pathways in the vicinity (∼3 cm radius) of the on-target location. The carotid bifurcation has a complex web of small nerve branches (Fjordbakk et al., 2019a); we traced the main nerve trunks (known to have motor branches) from their closest point at the stimulation location to their terminal muscles. The main nerve trunks identified in the microdissections were the hypoglossal nerve, vagus nerve, accessory nerve, superior laryngeal nerve, and recurrent laryngeal nerve, which branches off the vagus nerve in the chest and courses cranially to the larynx (Supplementary Figure 2-4). The microdissection showed the SLN pathway was closer to the stimulation location (∼1-1.5 cm) compared to the hypoglossal and accessory nerves (∼2-3 cm, although the distance was dependent on the surgical pocket size and the degree of retraction). Based on distances from the stimulation location, we anticipated that the SLN would be activated at lower thresholds as compared to the other two nerves. The activation thresholds of these nerves were quantified based on evoked responses from their target muscles in subsequent experiments.

### Identification of muscle groups activated during BAT stimulation

In our pilot study, we visually identified muscles that were activated during BAT. These muscles were instrumented with EMG electrodes in subsequent animals to measure dose-response curves during BAT. Figure 4 shows identified off-target neck muscle groups and their relative location in the surgical pocket. The main identified muscle groups were the CA, the CT, the TN, the CS, and the SC, where the SC is the swine equivalent of the sternocleidomastoid in humans. The average activation thresholds across the cohort (n=8) are reported in Figure 4C and Table 1 below. The CS had the lowest activation threshold, followed by the CT, the TN, the CA, and the SC. All muscle activation thresholds were comparable to stimulation amplitudes (∼3 mA) required to cause initial decrease in BP (Figure 3C).

**Figure 4:**
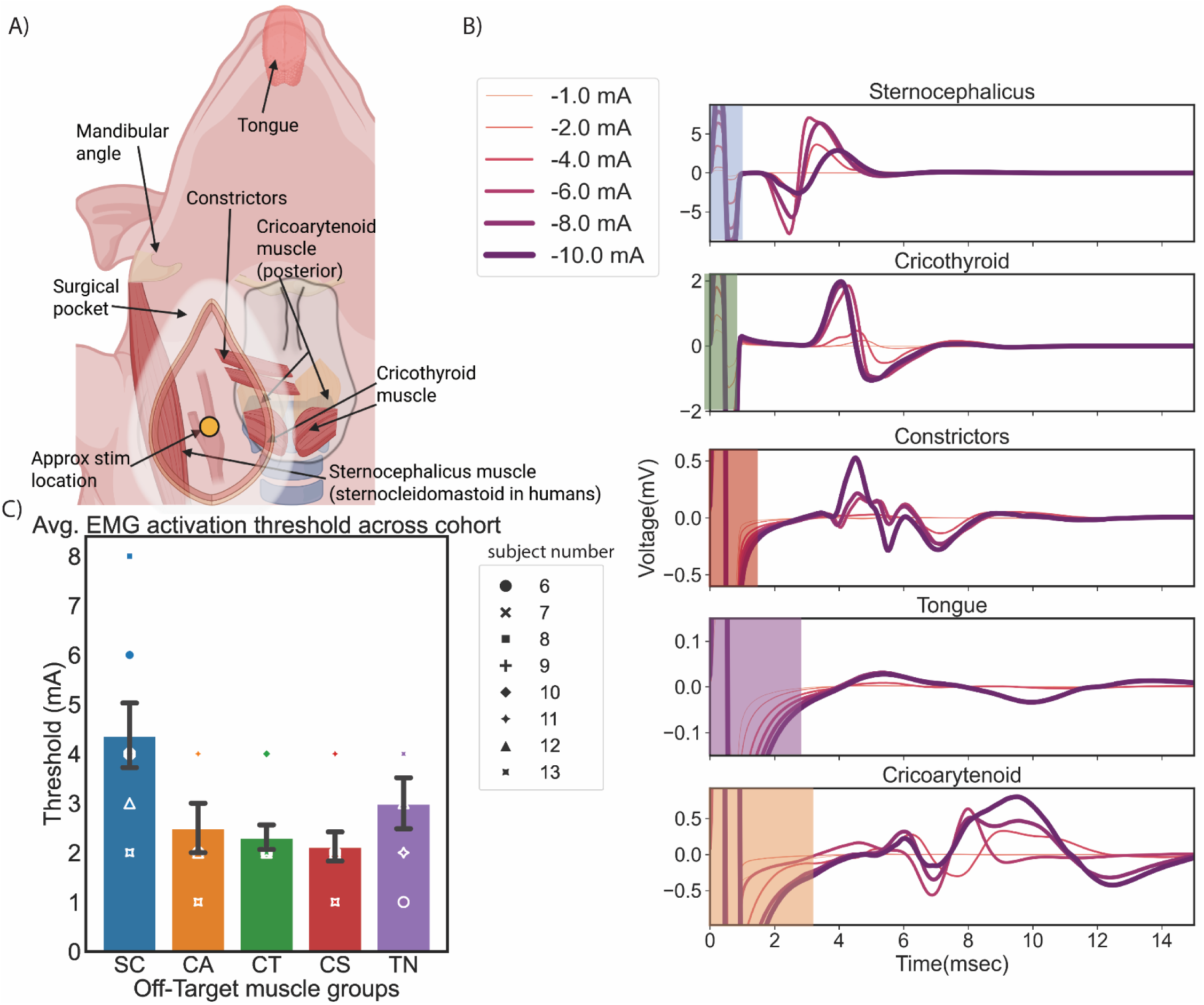
Identification of off-target muscle groups. A) Diagram of identified swine muscle groups that were activated during stimulation at the approximate on-target location indicated by the yellow circle. B) Recorded EMG from activated muscle groups across stimulation amplitudes for a representative swine. Increasing line thickness denotes increasing stimulation amplitude (1-10 mA). Highlighted color bands denote stimulation artifacts in the recorded signal, which were omitted during AUC calculation. C) Activation thresholds (mean ± SE) for off-target muscle groups across swine (n=8). Thresholds were visually identified as the lowest stimulation amplitude in the dose-response curve (with ∼1 to 2 mA step size) that evoked EMG responses for that muscle group (see Table 1 and Supplemental Section “Cohort EMG recordings for selecting activation threshold per muscle”). SC – Sternocephalicus, CA – Cricoarytenoid, CT – Cricothyroid, TN – Tongue, CS – Constrictors.

**Table 1:** Average muscle activation threshold. (the stimulation amplitude at which the first evoked muscle activity was recorded) per muscle group across cohort (n=8).

| Muscle group | Average activation threshold (mA): mean ± SD |
| --- | --- |
| Constrictors (CS) | 2.12 ± 0.83 |
| Cricothyroid (CT) | 2.31 ± 0.70 |
| Cricoarytenoid (CA) | 2.5 ± 1.22 |
| Tongue (TN) | 3.00 ± 1.26 |
| Sternocephalicus (SC) | 4.37 ± 1.84 |

The latency and activation thresholds of the EMG responses varied between muscle groups (Figure 4B), consistent with variable distances from the nerve pathways coursing near the stimulation location to the terminal muscles. Visually identified muscles activated during BAT and the latency of the response were used to inform subsequent transections to identify presumptive responsible neural pathways in the vicinity of carotid bifurcation.

### Functional verification of off-target nerve pathways using sequential transections

Microdissections from pilot swine and prior literature were used to identify segments of off-target nerves running proximal to the carotid bifurcation innervating activated muscle groups (CA, CT, TN, SC, and the CS). The CA and the CT are innervated by the branches of the SLN (Settell et al., 2020), the TN is innervated by the hypoglossal nerve (Shcherbatyy et al., 2008), and the accessory nerve innervates the SC (Johal et al., 2019)(Figure 5A and B). EMG responses at a subset of stimulation amplitudes were compared before and after transection to confirm the nerves responsible for muscle activation.

**Figure 5:**
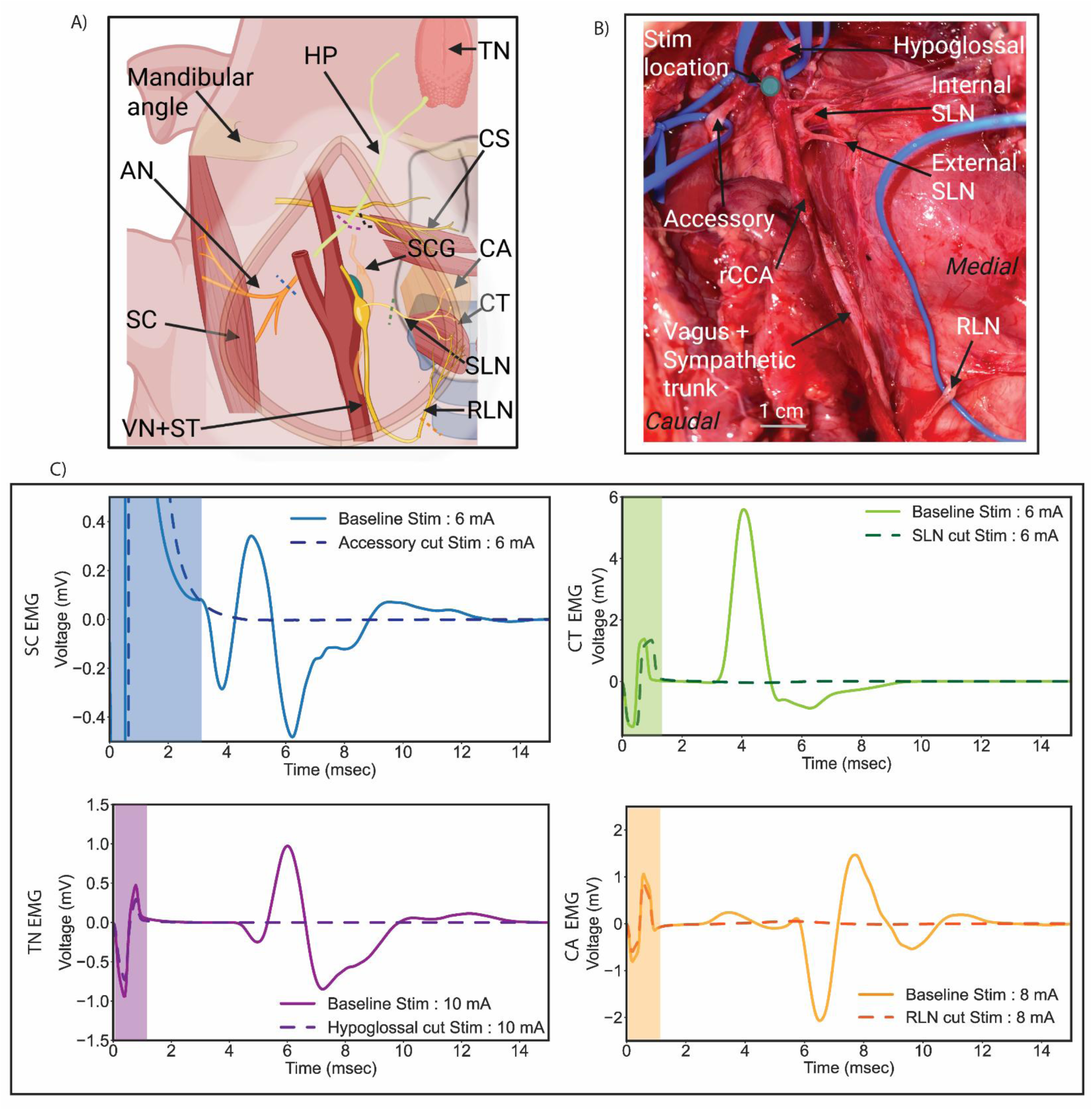
Identification of off-target nerve pathways. A) Diagram (not to scale) and B) in vivo photo of identified off-target nerve pathways with their respective muscle groups. C) Recorded evoked muscle response (EMG) from activated muscle groups during baseline and after nerve transection. Shaded areas represent stimulation artifact. Additional cohort data in Supplemental section “Additional swine nerve transection data”. SC – Sternocephalicus (sternocleidomastoid in humans), CA – Cricoarytenoid, CT – Cricothyroid, TN – Tongue, CS – Constrictors.

Representative EMG traces from muscle groups before and after their identified off-target nerve transections are shown in Figure 5C; additional data from other swine are reported in supplemental section *Additional swine nerve transection data*. As anticipated, transections of the RLN near its insertion to the CA eliminated the CA EMG response, and transections of the SLN as it branches from the nodose ganglion eliminated the EMG response recorded from the CT muscle. Transection of the hypoglossal nerve trunk as it coursed dorsal towards the foramen near the carotid bifurcation eliminated the EMG response of the tongue. The SC muscle EMG response was eliminated after transection of the accessory nerve trunk isolated near the carotid bifurcation.

Identifying the off-target nerve trunk responsible for the activation of the CS muscle was less straightforward than the other muscle groups. While transecting the RLN and the SLN reduced the evoked muscle signal, it did not eliminate it completely. Transection of additional small branches from the glossopharyngeal plexus was necessary to eliminate the EMG signal completely (Supplementary Figure 6).

### Dose-response relationship for identifying the therapeutic window with on- vs off-target thresholds

BAT efficacy is limited by side-effects of neck and jaw pain, cough, and breathing issues, putatively caused by activation of off-target neck muscles. All off-target muscles were activated in a dose-dependent manner across the cohort (Figure 6A). The stimulation dose-dependent responses between BP decreases and neck muscle EMG signals were compared (such that maximum response for both the EMG and BP reduction were equated to 1, see Methods section “Data Analysis”) across the cohort. The cricothyroid and constrictor muscles consistently had the lower thresholds for activation across the cohort. At the amplitudes tested, the threshold for reduction in BP coincided with the threshold for activation of these two muscles groups (Figure 6B), suggesting that it was not possible to elicit a BP response through BAT without concurrent muscle activation. Lastly, the CS AUC data (muscle group with the lowest threshold) versus the normalized BP reduction values were plotted to understand what percentage of off-target activation was evoked for a given reduction in BP, to understand if there were instances where it was possible to get greater reduction in the BP as compared to muscle responses. A linear relationship along the unity line between the two variables (gray dotted line, Figure 6C) would indicate that there was equivalent dose responsiveness to on and off-target activation. However, the majority of the datapoints lay above the unity line, which indicates larger muscle responses compared to decrease in BP at a given stimulation amplitude. For example, ∼50% of maximum possible BP reduction yielded a ∼65% of maximum evoked muscle response. A few stimulation amplitudes that resulted in the largest BP reduction evoked ∼80% of total recorded muscle activation. For better therapy efficacy, the goal would be to have most datapoints in the lower right half of the plot.

**Figure 6:**
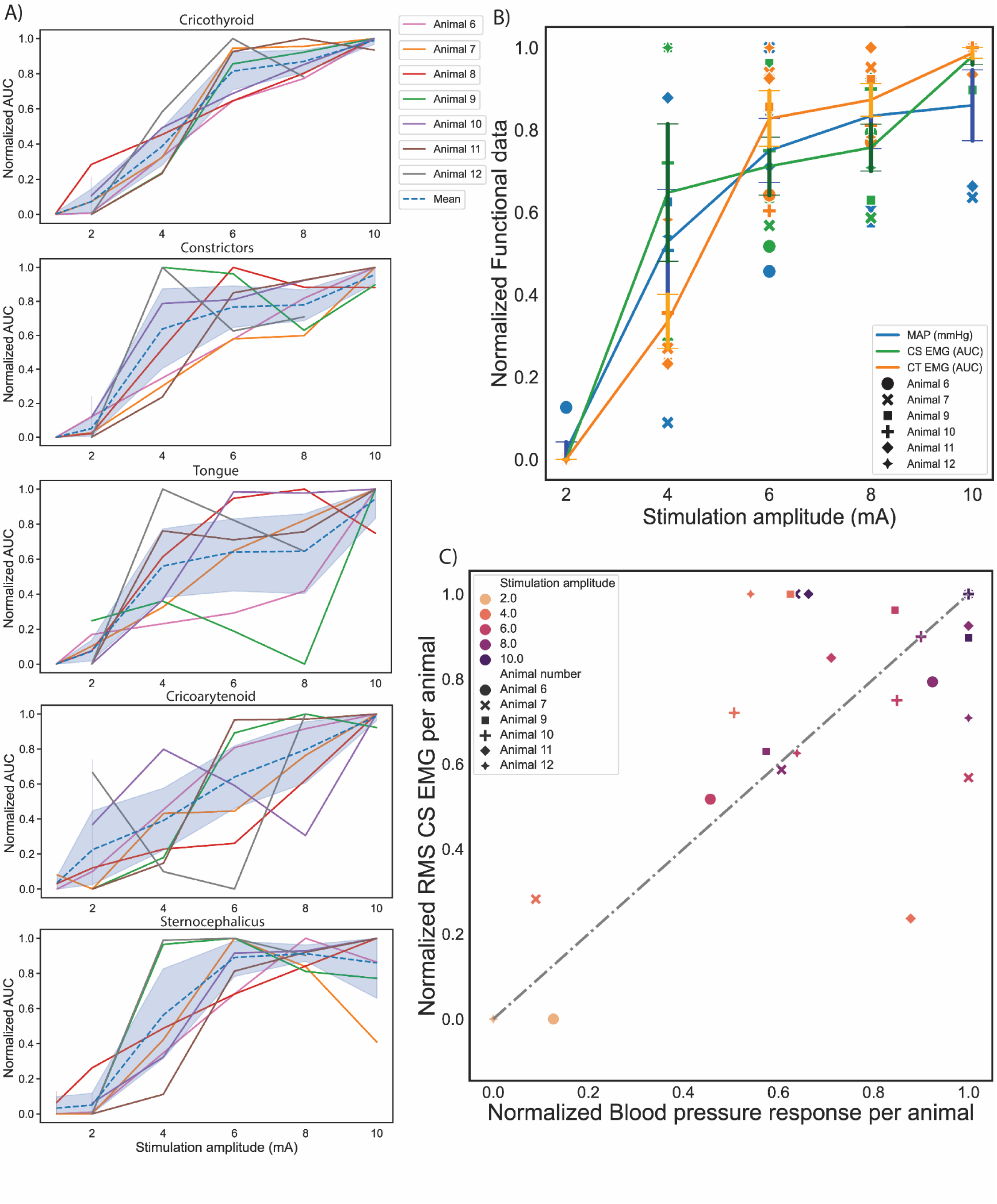
Stimulation dose-dependent responses of on- vs off-target activation. A) Normalized area under curve (AUC) for each muscle (EMG) group per swine. The blue line for each plot shows the mean and the shading shows the 95% confidence interval. B) Mean ± standard error of dose-response of normalized BP reduction (on-target, blue) and AUC of evoked EMG response of muscle groups with lowest activation thresholds (off-target) across the cohort (constrictors, green line and cricothyroid, orange line). Comparison of responses highlight that across all stimulation amplitudes tested; evoked muscle responses have a higher activation than decreases in BP. Individual swine data represented for every stimulation amplitude. C) Normalized mean CS EMG response per swine versus normalized BP responses per swine. Swine 13 was omitted from analysis due to atypical EMG response (see Supplemental Figure 9). The CS was chosen for analysis as the muscle group with the lowest activation threshold.

## DISCUSSION

BAT is used clinically to treat drug-resistant hypertension and heart failure with reduced ejection fraction. The device, once implanted, undergoes a series of dose titrations (increases in stimulation amplitude) over 3 months to reach the highest tolerable dose. However, therapeutic efficacy across large studies is severely limited, and the stimulation levels are variable across patients. The dosing is largely driven by individual tolerance of side-effects instead of consistent on-target engagement of baroreceptors (Schmidt et al., 2020). Multiple studies have suggested that these side-effects are largely due to the unwanted activation of neck muscles (Nicolai et al., 2020; Tosato et al., 2007; Yoo et al., 2013). Herein, we identify possible neuroanatomical substrates that are responsible for therapy-limiting side-effects in a swine model and their respective activation thresholds. As the swine model is translationally relevant to clinical therapy deployment (Lunney et al., 2021), the findings herein can be used as a foundation for electrode design testing, computational modeling of stimulation parameters, and exploring the anatomical framework for other cranial nerve therapies.

### Anatomy and physiology considerations for BAT clinical translation

#### Comparison of swine and human carotid anatomy

For BAT electrode design, it is important to understand the anatomy of the swine model in relation to the human. The swine CCA is slightly smaller in diameter (4-6 mm) than the human CCA (6-9 mm) and more closely comparable to the human ICA diameter (Yin et al., 2024). By comparison, the commonly used canine model CCA is notably smaller (3.7 ± 0.23 mm) (Martínez Moreno et al., 2019). From a surgical perspective, the swine CCA is located ∼45-75 mm from the skin which is ∼2x deeper than in humans (17-30 mm). The human carotid bifurcation is traditionally where the CCA splits into the ECA and the ICA structure. From caudal to cranial, from the carotid bifurcation, the human ECA further branches into the LA, the FA, the OA, the APA, and the MA with some inter variability (Devadas et al., 2018). In some humans, the SLA branches off the ECA caudal to the LA as compared to swine wherein it branches off the CCA (Settell et al., 2020). The nomenclature and branching pattern are important to identify the correct “on-target” BAT bifurcation location.

BAT is delivered at the carotid bifurcation, defined as where the CCA splits cranially into the ECA and the ICA in humans and into the ECA and the APA in pigs, but the APA nomenclature is debated. The main anatomical difference between humans and swine, as it pertains to BAT, is the contested and varied nomenclature of the on-target artery (ICA in humans) and its termination point. The majority of swine literature reports that the ICA is present at the tympanic cavity and is “replaced” by the APA at the CCA-ECA carotid bifurcation (Godynicki & Frackowiak, 1979; Iosif et al., 2016; McGrath, 1977; Reinert et al., 2005; Tandler, 1906), while some report the ICA as being present at the bifurcation (Graczyk et al., n.d.; Schummer et al., 1981), especially in reference to swine BAT models (Bushi et al., 2008; Fjordbakk et al., 2019a; Palmisano et al., 1990; Strauss et al., 2017). However regardless of naming differences, the arterial branch in question terminates in the RT, a highly vascular network of small arteries, which exists in sheep, cats, goats, oxen, and pigs (Arakawa et al., 2007; J. Edwards et al., 2022; Graczyk et al., n.d.; McGrath, 1977; Strauss et al., 2017) but not in humans (De Gutiérrez-Mahoney & Schechter, 1972; Minagi & Newton, 1966; Ota, 2023). In humans, the ICA (through its own thermoregulation system) directly supplies blood to the brain, while in swine, the RT thermoregulates blood supply to the brain to help protect cortical tissues from damage (Hayward & Baker, 1969).

For successful BAT translational models, the focus is not just the nomenclature of the arterial structure for the on-target location, but the physiological implications due to differences in anatomy between the two models as elaborated below.

#### Comparison of swine and human baroreflex physiology

Our data suggest that the on-target artery, whether labelled as the ICA or APA in swine, caused repeatable decrease in BP in a location similar to humans (Fjordbakk et al., 2019a; Linz et al., 2013, 2016). The decrease in BP for a given stimulus intensity is driven by baroreflex sensitivity (BRS) or baroreceptor-HR reflex (Parati et al., 1988). BRS is the quantification of how changes in detected BP (afferent signaling measured at baroreceptors) translates to parasympathetic/sympathetic signaling changes (efferent signaling measured in HR) (La Rovere et al., 2008). BRS is commonly quantified as the slope of the line or rate of change in R-R interval (ms) per change in systolic BP (mmHg). It is a clinical metric used to understand the severity of hypertensive patient pathophysiology and thus an important comparison between humans and swine for clinical translation of bioelectronic therapy. Swine have a BRS that is less sensitive but is closer to humans than in canines (which are highly sensitive) (Booth et al., 1960; Rigel & Millard, 1992; Slinker et al., 1982), yet historically canines have been the most commonly used preclinical BAT model.

BP changes can also be driven through activation of the afferents innervating the baroreceptors instead of the baroreceptors themselves in the arterial wall. These afferent fibers form the carotid sinus nerve (CSN). The swine CSN consists of large diameter myelinated A fibers (type I baroreceptors maintaining BP during sudden changes), small diameter A fibers, and unmyelinated C fibers (type II baroreceptors regulate basal BP) (Seagard et al., 1990, 1993; Suarez-Roca et al., 2021). The composition of the CSN in swine closely mimics that of humans in terms of nerve diameter, fiber diameters, and anatomical complexity (Fjordbakk et al., 2019b). Thus, the activation thresholds for on-target engagement of afferent baroreceptor fibers between the two models would be comparable. Our data suggests that a 1 mm disk at the on-target location requires ∼4-6 mA to drive appreciable reduction in BP in anesthetized swine. These stimulation amplitudes are comparable to clinically reported values at the end of 3 months titration window (from 4.5±2.5 mA to 6.8±2.4 mA) for the new Neo design (2 mm disk electrode) with maximally tolerated side effects (Abraham et al., 2015a). These data suggest that the on-target engagement between the two models could be comparable and warrants a comparison between their off-target pathways.

#### Comparison of swine and human off-target nerves

Our study identified off-target nerves (Figure 7 and Figure 8) at a 0.8-3 cm distance from the on-target location, for which the pig BAT activation thresholds are similar or lower to those required to evoke decrease in BP. All these nerves are reported to be at comparable distances and orientation in humans (Pinto et al., 2024). The fascicular organization and epineural thickness of swine and human cranial nerves are closer than those in canines, which have a thicker epineurium so a higher activation threshold (Pelot et al., 2020; Settell et al., 2020; Yoo et al., 2013). As canines have a higher baroreflex sensitivity with a higher threshold for off-target activation as compared to humans, it could be possible that previously reported BAT preclinical research in canines (Iliescu et al., 2014; Zucker et al., 2007) may have overestimated the effect of BAT, while underestimating the amount of off-target activation. The similarity of off-target nerves and BAT responses between swine and humans underlies the translational relevance of the framework reported in this paper.

**Figure 7:**
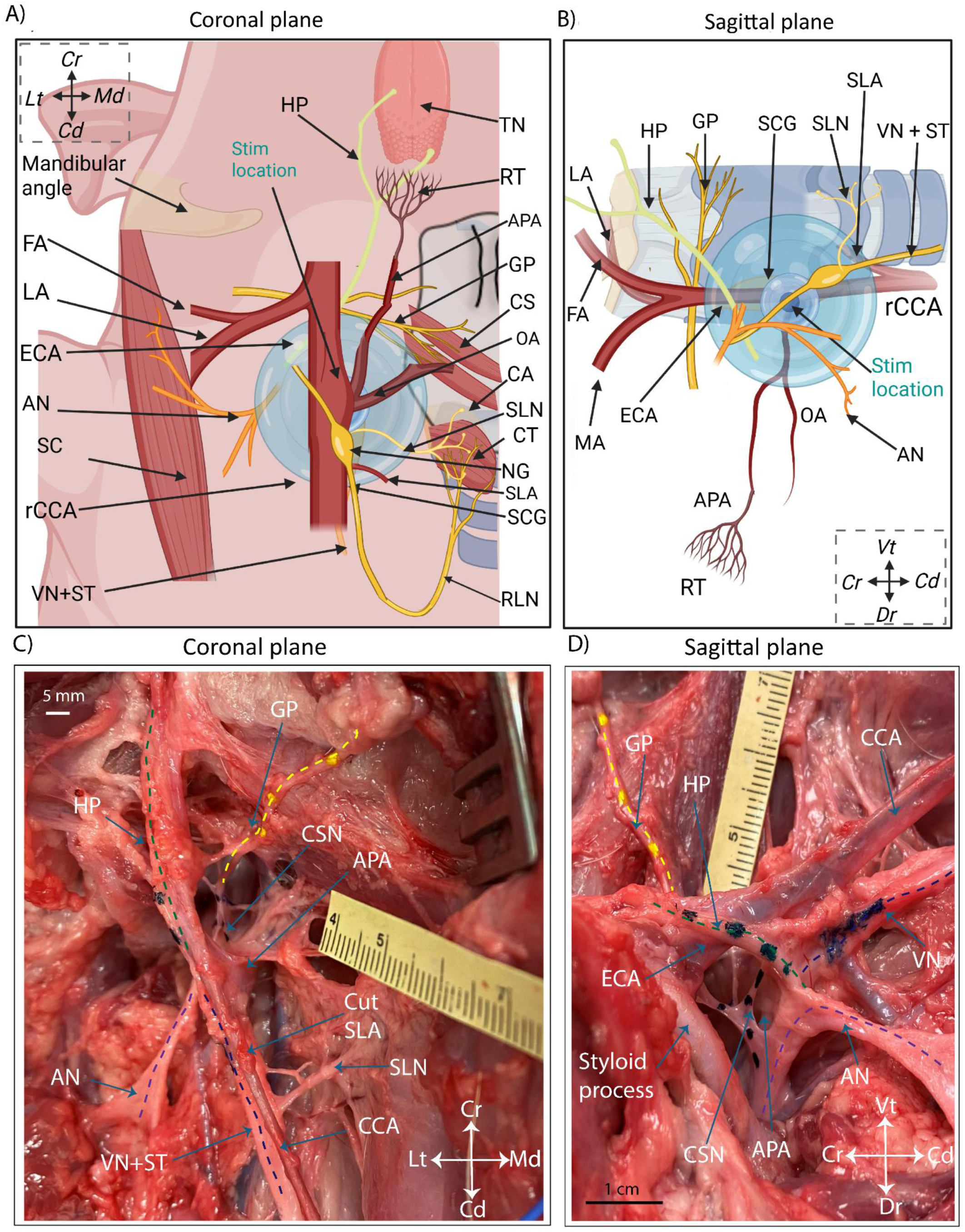
Summary figure identifying off-target muscles and nerves in BAT. A) Ventral view of swine carotid bifurcation anatomy and identified off-target muscles groups with their respective off-target nerve pathways. The representation highlights medial-lateral spatial distribution of the off-target nerves surrounding the stimulation electrode location and estimated volume of activation (blue bubble with color gradient indicating fall-off of electric potential with distance [dark (high) - light (low)]). B) Sagittal view of swine carotid bifurcation anatomy showing the ventral-dorsal spatial distribution of the off-target nerves. C) Coronal photo of cadaver microdissection showing the GP nerve (yellow dotted line) with multiple nerve branches as it travels dorsal towards the APA notch. D) Sagittal view of the GP nerve as traverses medial-lateral access with a nerve branch (possible CSN) innervating the APA-ECA bifurcation notch on the dorsal side (black dotted line). Off-target nerves, the VN (dark blue dotted line), the AN (purple dotted line) and the HP (green dotted line) as they travel dorsal to enter the jugular foramen.

**Figure 8:**
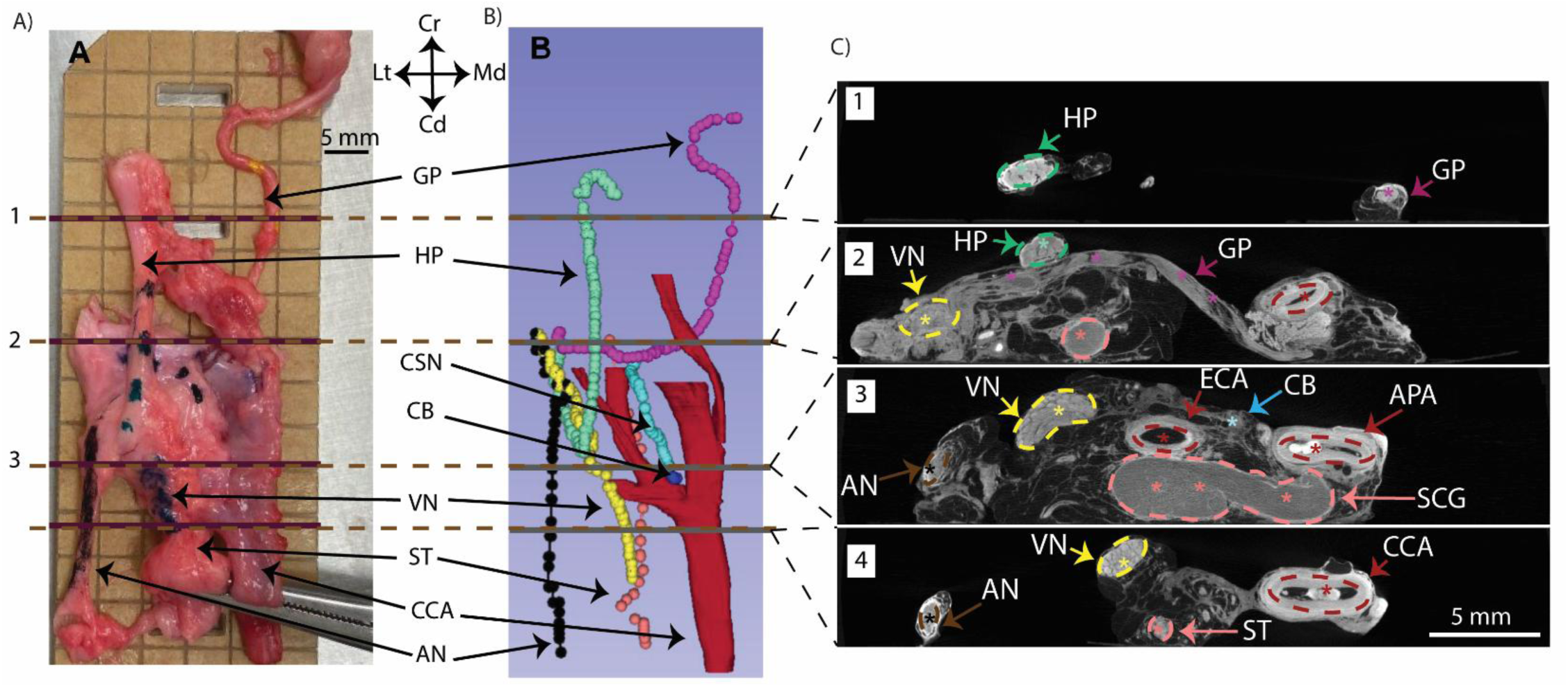
Imaging showing proximity of on-target location with off-target nerves in BAT. A) *En bloc* dissected sample fixed in 10% neutral-buffered formalin (NBF) and 70% ethanol was arranged on acrylic while maintaining approximate *in vivo* orientation for microCT imaging. B) MicroCT imaging and 3D volume rendering of sample (A) to visualize 3D arterial and nerve anatomy. The CSN (blue) branching off the GP (magenta) and innervating the identified CB (dark blue). C) Cross sections demarcated in panels A and B to visualize nerve sizes, fascicular organization, and on- vs. off-target distances. The cross section in subpanel 3 shows the CB and hence the approximate stimulation location at the APA. The off-target nerves are all 1 - 2.5 cm radius away from the stimulation location.

#### Loss of coordinated activity between activated muscles during tasks

Although the off-target nerves have multiple fascicles, the biggest driver of differences in activation thresholds is the distance from BAT electrode placement to the nerve trunk itself. The differences in distance to each fascicle within that nerve trunk is comparatively much smaller. This would mean that the difference in current threshold for getting any activation of motor fibers within the nerve trunk threshold and all of motor fibers in that nerve trunk would be relatively small. Thus, all equal diameter motor fibers in an off-target nerve would be activated synchronously (at the same instant if at the same distance).

However, a single off-target pathway can have multiple motor fascicles firing in varying coordinated patterns controlling different muscles. For example, the hypoglossal nerve is known to innervate 3 different muscles of the tongue (genioglossus: draws the tongue forward from the root, hyoglossus: retracts the tongue, and styloglossus: depresses its side) and has branches to the omohyoid, sternohyoid, and sternothyroid muscles (Lin & Barkhaus, 2009). Synchronized activation of all motor fibers could offset the coordinated cascade of signaling required between different muscle groups for certain tasks such as swallowing or speaking.

#### Activation of neck muscles can change breathing and the HR/BP baseline

Patients may habituate and become desensitized to reported unwanted side-effects such as hoarse voice, jaw pain, etc., over long duration titration windows (∼3 months) to reach the intended therapy intensity (Fisher et al., 2021). Even though these symptoms can be managed in the tolerable range, the effect of activated muscles and their off-target nerves on long-term BAT efficacy could theoretically go beyond tolerable muscle activation.

Our data shows that neck muscles such as the CT and the CS can be activated at lower thresholds than the threshold for decrease in BP. CT is responsible for opening vocal folds, which allows airflow during breathing. SC (sternocleidomastoid in humans) elevates the sternum and the 1^st^ two ribs, which helps the lungs expand during inspiration. Thus, activation of these off-target neck muscles over sustained periods of time may also cause changes in breathing patterns. Stimulating neck muscle proprioceptive sensory afferents alters cardiorespiratory brainstem neural circuits (Edwards et al., 2015). These fibers have slightly higher activation threshold than motor fibers, which innervate the identified neck muscles. Activation of these muscles could also cause closing of vocal folds that may alter central respiratory patterns due to sensory mediated reflexes. The changing patterns of the central respiratory drive and cardiovascular functions (especially since perfusion to organs and BP is tightly coupled (Meng, 2021)) could putatively cause drifts in BP baseline in a patient population suffering from hypertension. These drifts in BP may increase the risk of cardiovascular disease in the target population for BAT (Stevens et al., 2016). This could theoretically be one of the contributing factors as to why BAT is considered safe but has not shown to reduce mortality or morbidity in heart failure patients (Blanco et al., 2023; Zile et al., 2024).

#### Mapping of reported side-effects to the identified off-target nerves

The off-target nerves identified herein were previously reported in clinical studies as having a chance of being damaged or activated during BAT surgical procedures (FDA, 2019). We categorized the clinically reported side-effects to their possible neural source in Table 2.

**Table 2:** Clinical reported side-effects and their possible off-target somatic low threshold motor nerves. SLN-superior laryngeal nerve, RLN-recurrent laryngeal nerve, AC - Accessory nerve, HP – Hypoglossal nerve, GP – Glossopharyngeal nerve, VN – Vagus nerve

| Clinical reported side effect | Possible off-target nerves | Reference |
| --- | --- | --- |
| altered sensory function of pharynx and oropharynx | VN, GP, SLN | (FDA, 2019; Steele & Miller, 2010) |
| facial palsy/paralysis | Cranial nerve VII | (Yang & Dalal, 2021) |
| altered sensory and motor function of tongue | HP, GP | (Snyder & Bartoshuk, 2016) |
| excessive salivation, dry cough | HP, VN, GP | (Al-Biltagi et al., 2022) |
| respiratory constriction and stertorous breathing | AC | (Andrew, 1955) |
| altered sense of taste | HP, VN, GP | (Heckmann et al., 2003; Witt & Reutter, 2015) |
| altered speech | HP, SLN | (Orestes & Chhetri, 2014) |

As continuous EMG readout from the responsible off-target muscle groups is impractical in humans, feedback about side-effects from patients during reprogramming stimulation programs at home could aid in selecting better stimulation contacts and programs. This concept of patient feedback for selecting programming parameters to reduce side effects is not new and is currently already in use in spinal cord stimulation, suggesting that its implementation in the clinic is a feasible goal (North et al., 2003).

#### Physiological effects of possibly activating the autonomic nervous system

This study mainly addresses large diameter motor somatic off-target fibers as they have the lowest activation threshold and are responsible for muscle activation contributing to reported therapy limiting side-effects. However, BAT for heart failure patients is predominantly a therapy of resetting the autonomic nervous system where the goal is to increase parasympathetic output while reducing sympathetic tone. BP and HR are tightly coupled and controlled to match varying physiological demands such as exercise. The VN, when activated (parasympathetic efferent B-fibers), causes bradycardia (drop in HR) and thus reduction in BP (Blanz et al., 2022). The sympathetic fibers when activated cause tachycardia (increase in HR) and thus an increase in BP. Previous publications and our pilot data reveal the superior cervical ganglion and the sympathetic trunk in the vicinity (∼5-10 mm) of the on-target stimulation location, in addition to the VN (Figure 8). This points to a likelihood that hypotension and hypertensive crises during BAT, both reported adverse events (Bakris et al., 2012; Bisognano et al., 2011), could be worsened due to irregular activation of these off-target pathways. Previous publications have also explored how activation of cervical sympathetic ganglia causes potent cerebral vasoconstriction (Kim et al., 2022). The sympathetic preganglionic B fibers and postganglionic C fibers are harder to activate than large diameter motor nerve fibers, so locating the electrode closer to baroreceptor afferents may make it more feasible to avoid the large diameter fibers. Future studies exploring the role of activation of autonomic nervous system components (VN, sympathetic ganglia, ST) towards BAT efficacy should be explored.

#### Complications in surgical cutdown for electrode implantation

Efficacy of BAT is highly dependent on precise placement of the stimulation electrode at the carotid bifurcation as indicated by clinical data (Weaver et al., 2016) and the results of this study. However, this targeting requires an invasive cutdown and clearing connective tissue in the bifurcation. This anatomical location has a dense mesh of afferents from baroreceptors and chemoreceptors, along with cross-connecting microbranches between the VN, the ST, the AC, the HP, and the CSN not visible without magnification (Figure 8) (Kikuta et al., 2019). Avoiding damage to these structures in a surgical implantation procedure along with limiting the time under anesthesia would be an improbable task and a major hurdle for broader adoption of BAT. Damaging the CSN would make the entire procedure ineffective while damage to the other cranial nerves may cause chronic issues such as Horner’s syndrome (damage to sympathetic trunk) and facial palsy (cranial nerve VII). These factors may help to explain why BAT clinical data with previous electrode designs (CVRx Rheos), that were wrapped around the CSN and required substantial dissection, had 24% unresolved procedural related adverse events. Moreover, the current clinical electrode (Neo) was designed to reduce the procedural event rate putatively due to less surgical dissection but may require more precise placement for effect.

### Implications for Neural Engineering

#### Electrode design and stimulation programming

The current BAT clinical electrode (Neo) was designed to address the aforementioned surgical issues associated with Rheos. Another key design change was to a monopolar electrode compared to the Rheos bipolar and tripolar configurations. This design change helped preserve battery life with comparable on-target engagement. However, the fall-off of the electric potential is 1/r for monopolar stimulation as compared to 1/r^2^ for bipolar stimulation, where r is the distance of the nerve from the electrode. Thus, the monopolar configuration is expected to activate a larger number of nerve fibers and at a farther distance as compared to bipolar stimulation. However, the current required to activate the on-target fibers would also be reduced during monopolar stimulation. Thus, it would be necessary to understand if both the intended and the off-target activation scales linearly between the two stimulation configurations.

For monopolar stimulation, using nerve distances from our swine framework and assuming the monopolar disk electrode is directly on top of Aδ on-target fibers (Table 2), for approximately 20% of Aδ fibers to be activated, 100% Aα fibers are activated at the same location (Aδ threshold is approximated as 5x Aα threshold, Table 2) (Hussain et al., 2024; Musselman et al., 2021).If the Aα is moved 5-10 mm away from the stimulation location, 20-10 % of all Aα fibers would be activated compared to 20% Aδ activation at the stimulation location (assuming the closest off-target nerve viz SLN is 5-10 mm away). Thus, both on- vs off-target fiber types would have similar activation thresholds even though Aα’s are much farther away, as reflected in our data. The degree of separation between activation of the two fiber types (Aα vs Aδ) can potentially be increased using shorter pulse widths (Anderson et al., 2020; Grill & Mortimer, 1996; Kuykendal et al., 2017; Reich et al., 2015).

**Table 2:** Estimated differences in activation of on- vs off-target nerves. . Representative off-target nerves, SLN (closest) and Accessory (furthest) as motor nerves (Aα) with activation thresholds reported in literature.

| Anatomy structure | On vs Off-target | Fiber type | Activation threshold of fiber type in literature | Distance of fibers from stimulation location (R) | Estimated activation threshold multiplier of | Activation threshold in |
| --- | --- | --- | --- | --- | --- | --- |

**Table 2: Estimated differences in activation of on- vs off-target nerves.** Representative off-target nerves, SLN (closest) and Accessory (furthest) as motor nerves (Aα ) with activation thresholds reported in literature.
|  |  |  | *(references in text) (mA)* | *estimated from microCT and dissection (mm)* | *Aα at the stim location (assuming all fibers are at the stim location)* | *functional data (mA) Ref: Figure 3C and 4C* |
| --- | --- | --- | --- | --- | --- | --- |
| **SLN nerve** | *Off* | *Aα* | *0.1 - 0.6* | *~8-12* | *1 x* | *2.31 ± 0.70* |
| **Accessory nerve** | *Off* | *Aα* | *0.1 - 0.6* | *~15-25* | *1 x* | *4.37 ± 1.84* |
| **Baroreceptor afferents** | *On* | *Aδ* | *1.5 - 4* | *~1-8* | *~ (2.5-7) x* | *4 - 6* |
| **Baroreceptor afferents** | *On* | *C* | *8 - 10* | *~1-8* | *~ (13-17) x* | *NA* |

### Limitations

Our study had several limitations with the most important being that the experiments were performed in anesthetized swine. Differences between swine and human anatomy may influence the translation of these findings to the clinic.

#### Anesthesia

Isoflurane and propofol interfere with afferent neural signaling involved in BAT. In BAT clinical trials [28 patients], general anesthesia affected the mapping at the carotid artery to find reduction in BP. This led to a change in anesthesia protocol to have the lowest general anesthesia possible while maintaining an anesthetic plane and regional carotid plexus block. This was shown to improve the outcomes of carotid mapping to generate BP and HR changes (Schmidt et al., 2020). Our experimental protocol also had similar limitations due to anesthetic drugs, including isoflurane, propofol, and fentanyl blunting the sensitivity of the baroreflex. Future studies should consider using drug combinations of urethane/chloralose to reduce the effect of anesthesia on baroreflex blunting.

#### Drift in baseline BP over time and quantification error

Instrumentation calibration errors for measuring physiological signals can overestimate or underestimate the on-target activation. Thus, it is important to set the right voltage range for BP signal acquisition, as standard 1 V sampling range in LabChart resulted in noisy BP recordings in swine 1-5. These data were unreliable for quantifying small decrease in BP. Additionally, the sampling voltage range is adjusted during catheter calibration and cannot be adjusted after the BP signals have been recorded in post-processing. This calibration error was corrected in the experimental protocol refinement stage before dose-response curves data collection.

The baseline BP drifts over the duration of the experiment due to the surgical cutdown procedure, anesthesia, and hemodynamic changes due to bodily fluids. We attempted to mitigate this issue in dose-response curves by collecting all the data within 15 min. Additionally, we calculated the BP reduction compared to baseline just before the start of that specific stimulation amplitude, thereby accounting for some baseline drift. However, the magnitude of BP reduction can vary depending on the baseline hemodynamic range and autonomic tone. This could lead to quantification errors for comparisons across swine; however, our data suggests these differences were small. Additional studies are required to study BAT sensitivity across different autonomic tones and baseline BP values.

#### Surgical cutdown for targeting baroreceptors

The surgical cutdown procedure for targeting baroreceptors is highly invasive. The methodology requires clearing connective tissue at the carotid bulb to suture the electrode in place. It is highly probable that some axons in the connective tissue were cut. This could alter the sensitivity of BAT BP responses for the tested stimulation amplitudes.

#### Open versus closed pocket for BAT muscle activation

In electrical stimulation, the current spread from the stimulation electrode is contingent on the conductive medium. There are clear differences in the activation profile of nerve fibers based on the medium between the stimulation point and the nerve (fat, blood, edema, muscles, bone, etc.). In our experimental preparation, the off-target nerves were isolated from the stimulation electrode due to air in the open surgical pocket with surrounding muscles having been retracted. Extra fluid buildup around the stimulation electrode was also regularly removed using surgical gauze. Opening the surgical pocket to visualize and surgically isolate the off-target nerves increases the distance between the nerves and the stimulation electrode as compared to closed surgical pocket preparation. It is possible that the off-target activation of muscles would be at lower stimulation thresholds than in an open pocket.

#### Off-target activation of additional muscle groups

The off-target muscle groups activated during BAT were the CA, CT, TN, CS, and the SC. The trapezius muscle was also activated, but it was not included due to difficulty in consistent instrumentation and reliable EMG recordings owing to its anatomical location and motion due to stimulation (Supplemental Figure 7). The EMG signals would sometimes have a bleed though signal with a slightly different latency. These would indicate activated muscles in the vicinity innervated by the same or different neural pathways. The transections of coalesced nerve trunks eliminated all stimulation-evoked EMG signals, suggesting that there were few missed additional nerve pathways. However, there would be multiple other possible muscle groups innervated by the identified off-target nerves through other small nerve branches not reported herein. These muscles could be further separated from the EMG recording location and hence not completely captured in our EMG signals from specific anatomical locations. It would not be practical or feasible to record signals from all muscles innervated by the identified off-target nerves.

#### Limited sample size for dosing

Due to practical time constraints for experimental execution, we were limited in the step size for stimulation dose-response curves. The current steps (1-2 mA) were selected based on our pilot data to show clear dose-dependent BP responses. These stimulation amplitudes were not designed to tease out the granularity of off-target muscle recruitment. It is highly likely that the actual thresholds for EMG activation are between the threshold we are reporting and the next lower amplitude tested that did not evoke a response, i.e., slightly lower than the reported thresholds. However, the main objective of this study was to identify the neural substrates responsible for off-target muscle activation as well as to quantify the activation of off-target muscles in relation to decrease in BP. Our dosing strategy was designed to answer these questions.

#### Statistics

All statistics and analyses were post-hoc, not pre-registered, and therefore should be considered exploratory. Additional future experiments with pre-registration are required as confirmatory studies.

## CONCLUSION

Drug-resistant hypertension remains one of the leading causes of death and a huge burden on the health care system with BAT as a promising device-based solution. However, the efficacy of BAT is limited by side-effects with long periods of therapy titration and habituation. Here, we identified and confirmed off-target neck muscles that are activated during BAT and their neural pathways responsible for activation. This neuroanatomical framework can be used as a foundation for designing electrodes and building computation models for testing various waveforms to preserve target engagement while mitigating off-target activation.

## Acknowledgements

This work was supported through NIH/NIBIB R01 EB033403 and NIH SPARC Program. We would like to thank the staff of University of Wisconsin’s Center for Biomedical Swine Research & Innovation (CSBRI), Translational Research Initiatives in Pathology Laboratory (TRIP) and Research Animal Resources and Compliance (RARC) anesthesiology and imaging unit for their operational and veterinary support during these experiments.

## List of common acronyms

CCA: common carotid artery
APA: ascending pharyngeal artery
OA: occipital artery
ECA: external carotid artery
RT: rete mirabile
CT: cricothyroid
CA: cricoarytenoid
CS: constrictors
TN: tongue
VN: vagus nerve
AC: accessory nerve
HP: hypoglossal nerve
GP: glossopharyngeal nerve
SLN: superior laryngeal nerve
CSN: carotid sinus nerve
ST: sympathetic trunk
EMG: Electromyography
MAP: mean arterial pressure
BP: blood pressure
HR: heart rate

