## Supplemental material for "Identification of low threshold off-target activation pathways during stimulation of carotid baroreceptor afferents in swine"

### Supplemental Document

#### Methods:

Settings for Mean Arterial pressure calculation from recording blood pressure.

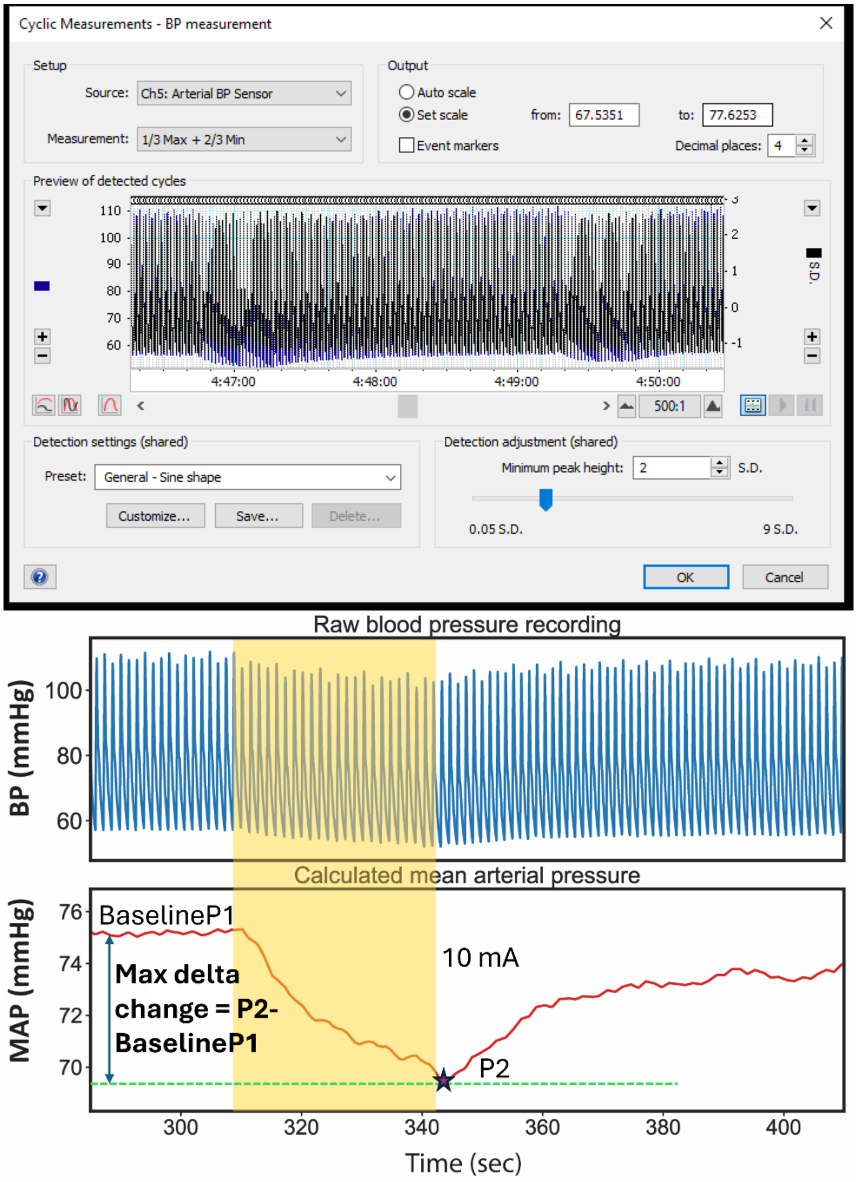


Supplemental Figure 1: Top: Settings for ADIntruments Labchart for thresholding raw blood pressure data to calculate Mean Arterial pressure. Bottom: BP response to stimulation calculated as Max delta change in MAP from baseline value.

#### Animal number table and stim params

| Subject number | Sex | Wt | Stimulation frequency (Hz) | Stimulation pulse width per phase (usec) |
|---|---|---|---|---|

| | | | | |
|---|---|---|---|---|
| Subject 1 | M | 35 | 75 | 350 |
| Subject 2 | F | 33 | | |
| Subject 3 | F | 40.3 | 25 | 250 |
| Subject 4 | F | 38 | 75 | 350 |
| Subject 5 | M | 40.1 | 25 | 350 |
| Subject 6 | M | 29.5 | 25 | 175 |
| Subject 7 | F | 41 | 25 | 350 |
| Subject 8 | M | 40.2 | 25 | 350 |
| Subject 9 | F | 38.3 | 25 | 350 |
| Subject 10 | M | 41.2 | 25 | 350 |
| Subject 11 | M | 32.5 | 25 | 350 |
| Subject 12 | M | 43 | 25 | 350 |
| Subject 13 | M | 33.8 | 25 | 350 |

Supplemental Table 1: Cohort subject details for weight, sex and stimulation parameters used for pilot and functional studies. Subject 1-5 had an old blood pressure calibration technique and were not included in dose response curve analysis.

#### Results:

##### Microdissection

Subject 3

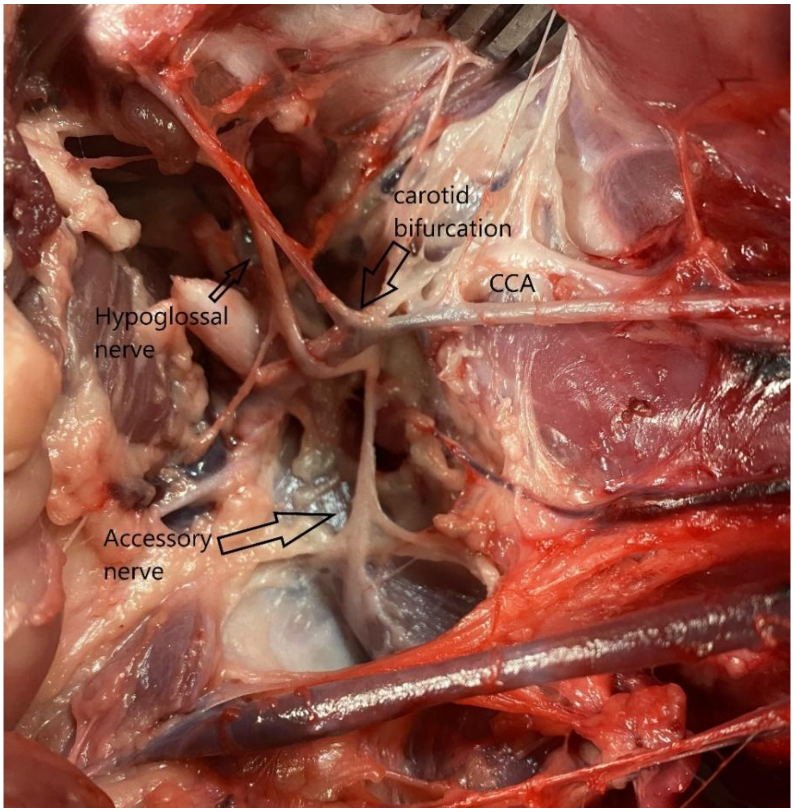


Supplemental Figure 2: Postmortem dissection to identify and trace off-target nerves hypoglossal and accessory back from their muscle groups to the carotid bifurcation. Both nerves enter the foramen dorsal to the bifurcation.

Subject 9

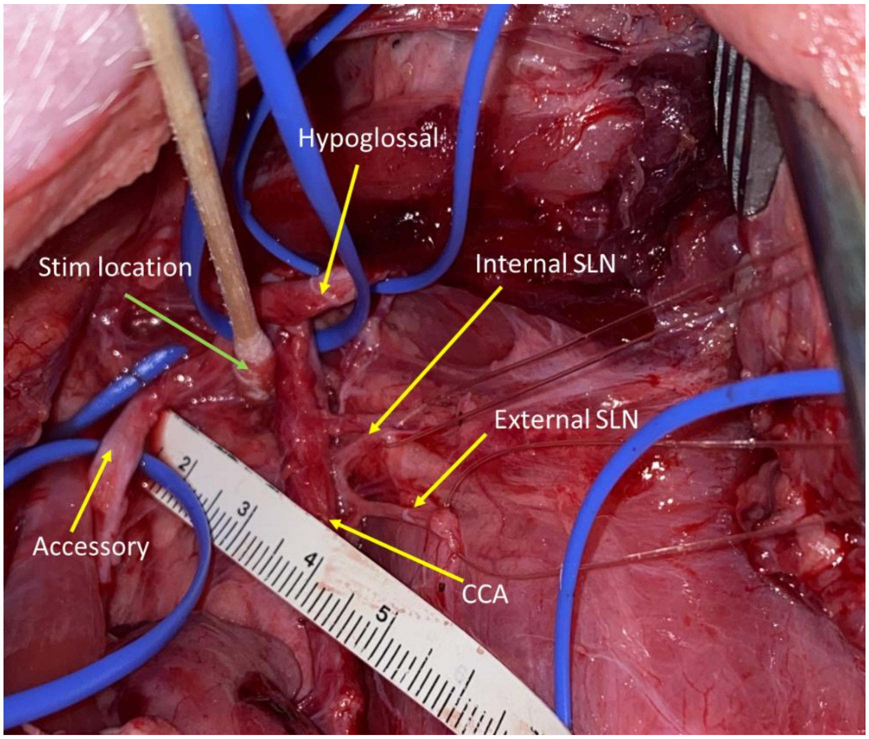

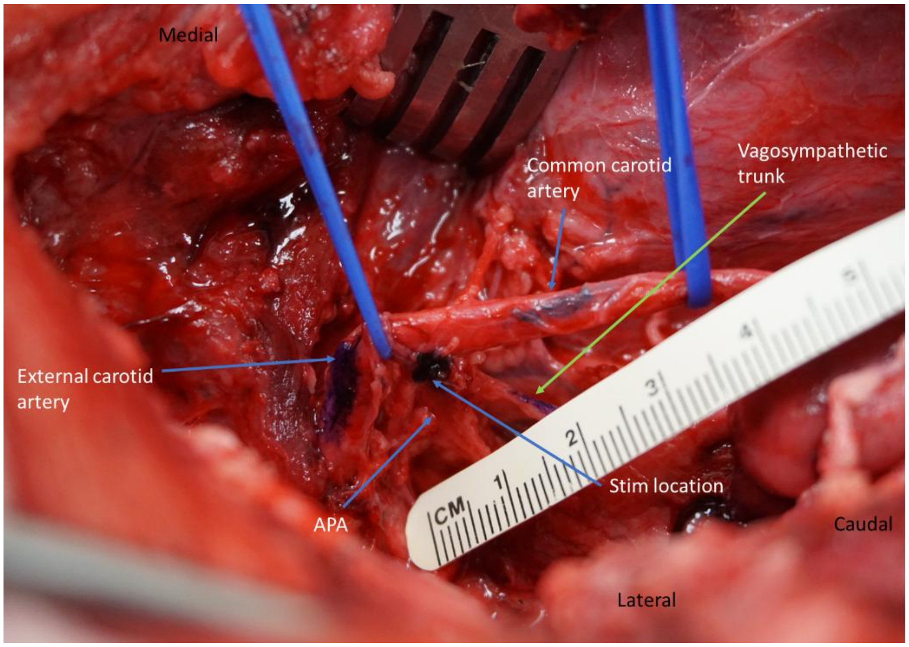


Supplemental Figure 3: Dissection showing off target nerves and their distances during stimulation (Top). (Bottom) Approximate stimulation location at the APA-OA bifurcation with the carotid artery retracted cranial and ventral.

Subject 13

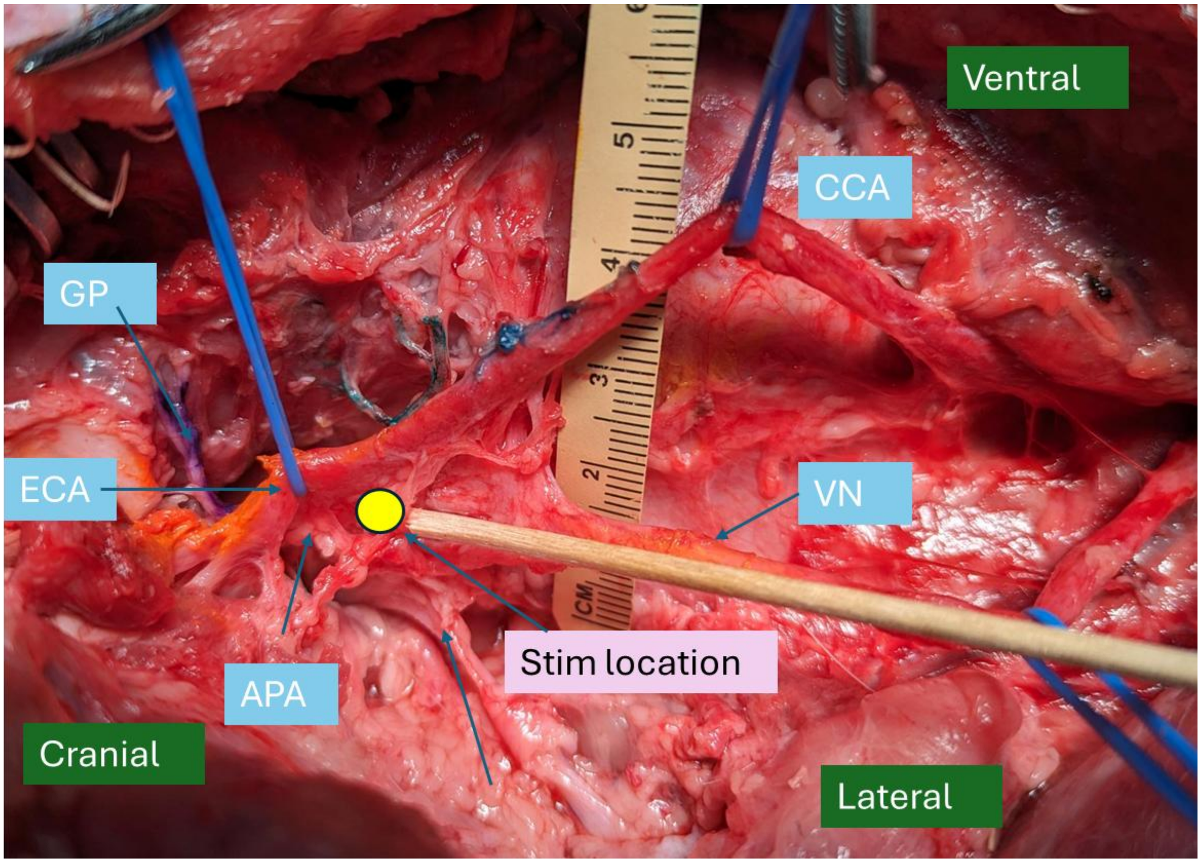

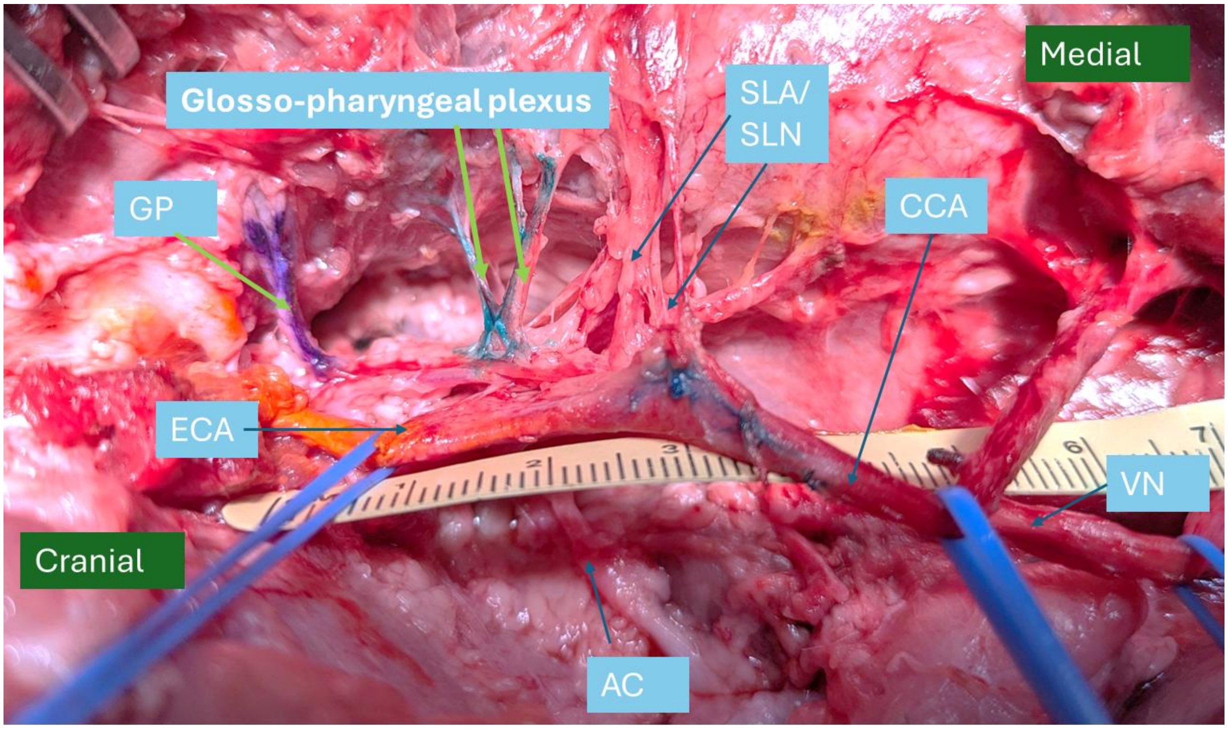


Supplemental Figure 4: Top) Post-mortem dissection showing approximate stimulation location at the APA-OA bifurcation with the carotid artery retracted medial and ventral. Bottom) Dissection showing glosso-pharyngeal plexus which innervates the constrictor muscle along with other off-target nerves. GP: glossopharyngeal nerve, AC – accessory nerve and VN vagus nerve.

#### Additional subjects nerve transection data

##### *Accessory nerve*

Subject 3

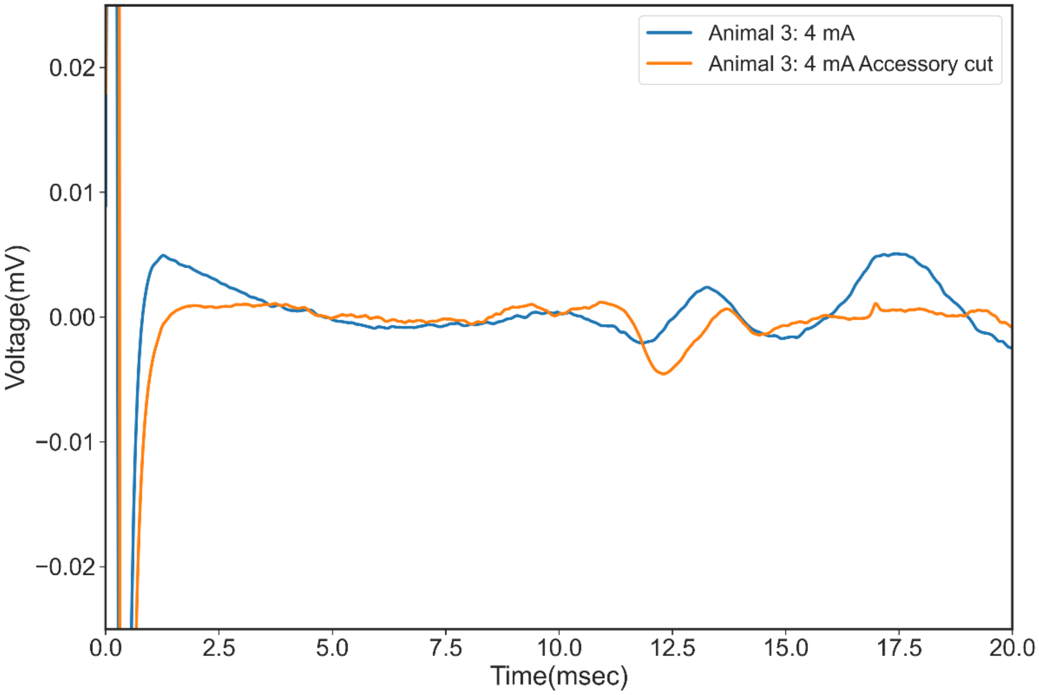

Supplemental Figure 4: Nerve transection of the only the external branch of the accessory nerve as it innervated closer to the muscle, eliminated only part of the EMG signal. This data led to a change in location of accessory nerve cut to be more cranial, medial/dorsal to the parotid gland exit point and closer to the stimulation location (as it traverses dorsal to the foramen) where the nerve is a well coalesced trunk with all its branches.

Subject 6

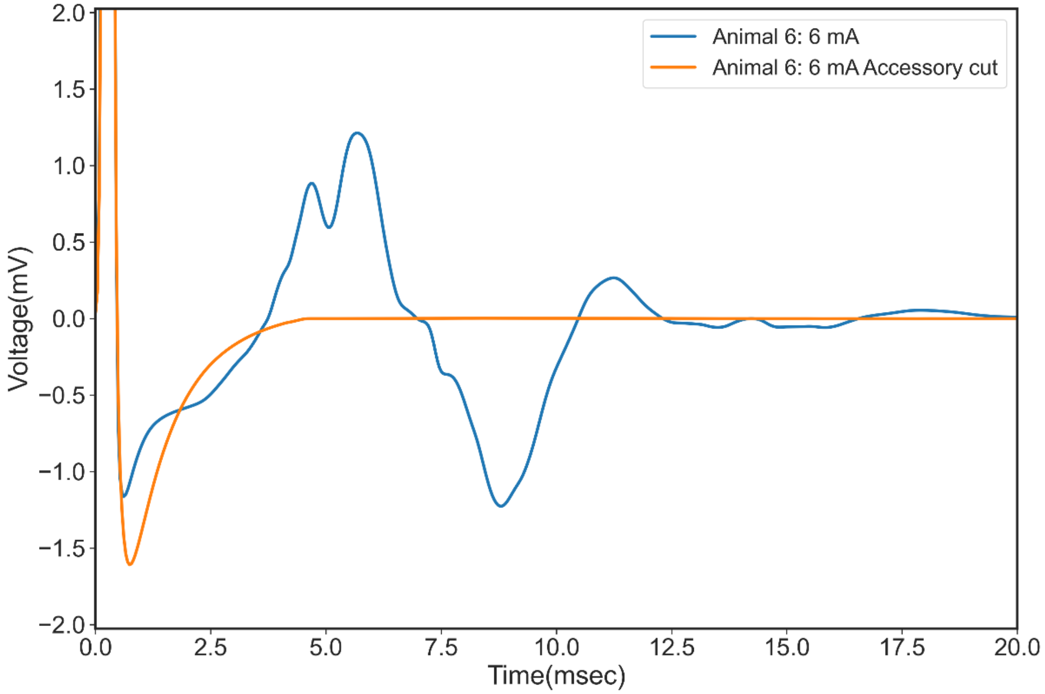


Supplemental Figure 5: Complete elimination of EMG post accessory nerve trunk transection

Subject 10

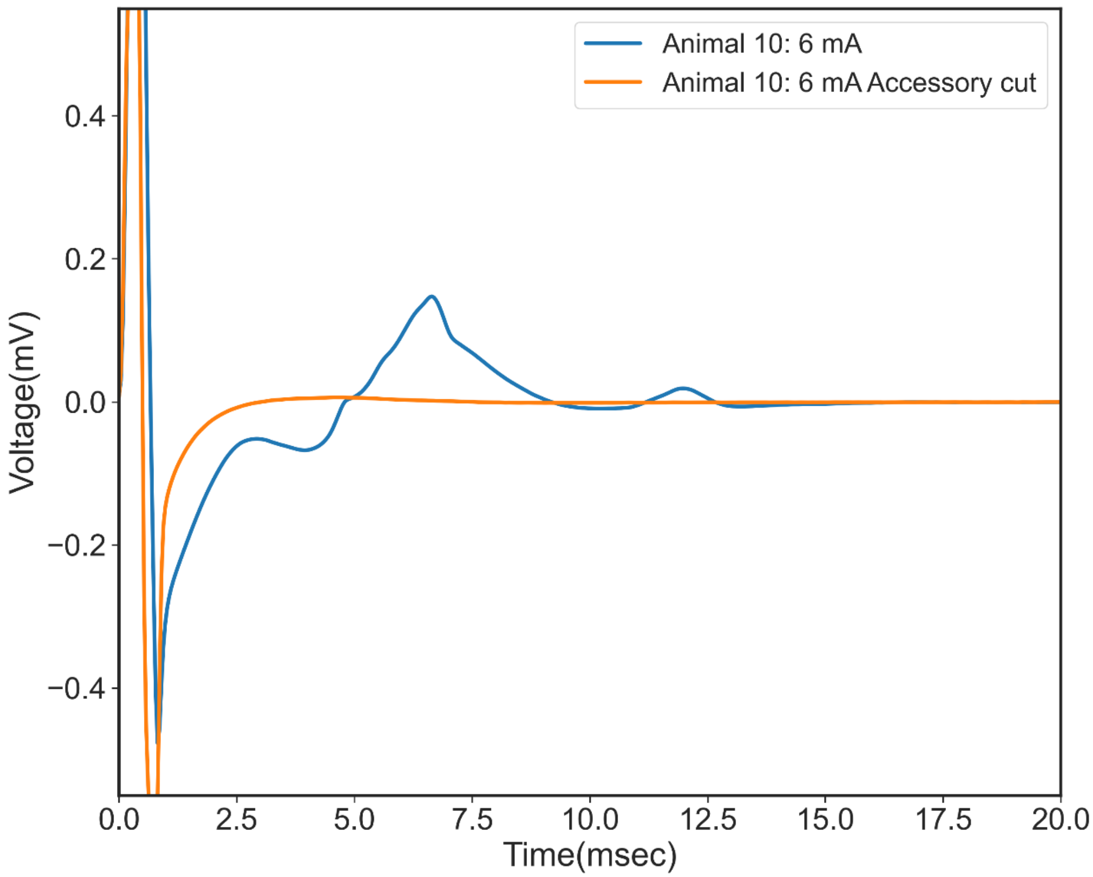


Subject 12

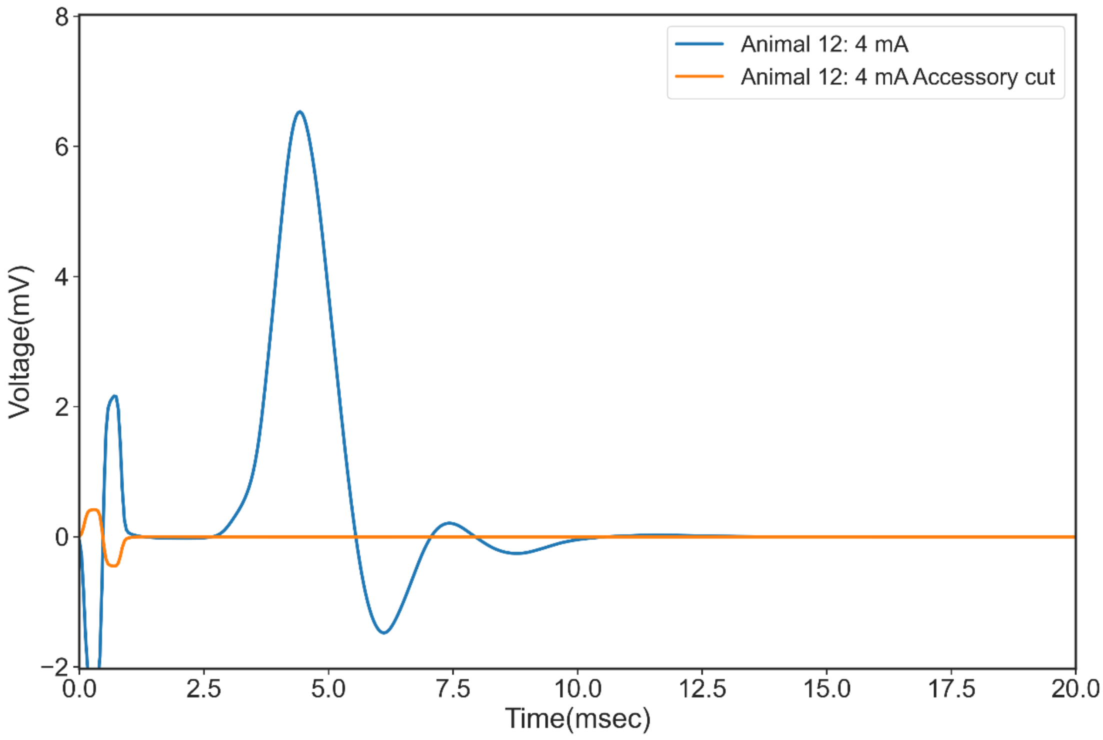


***Hypoglossal***

Subject 5

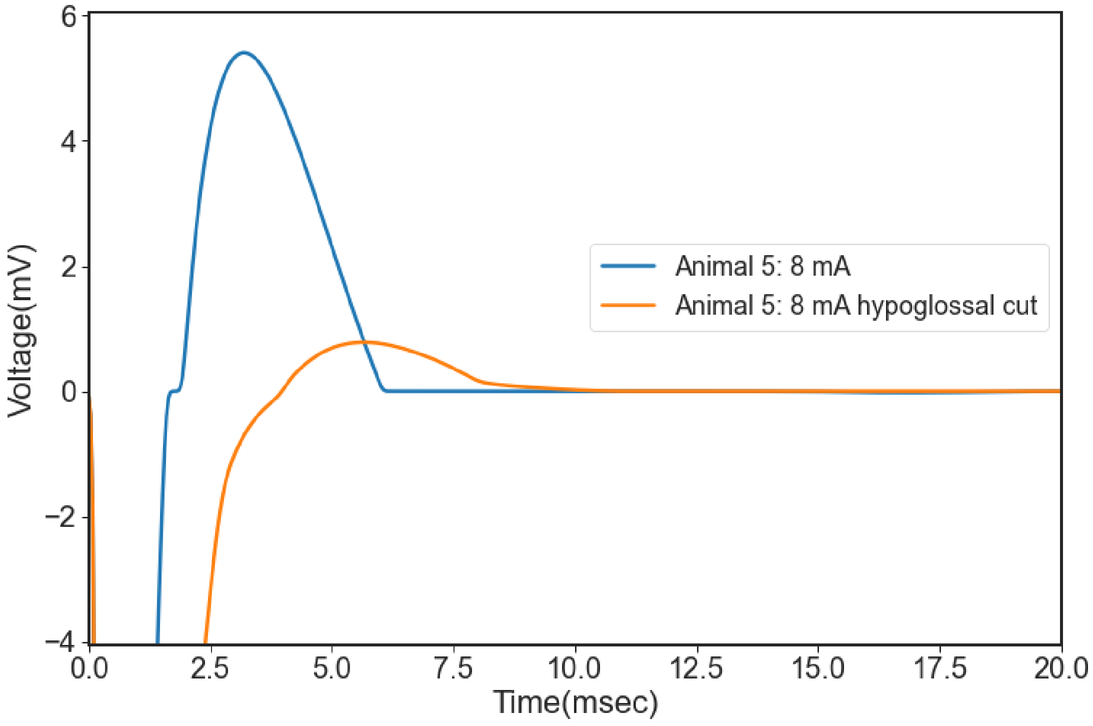


Subject 6

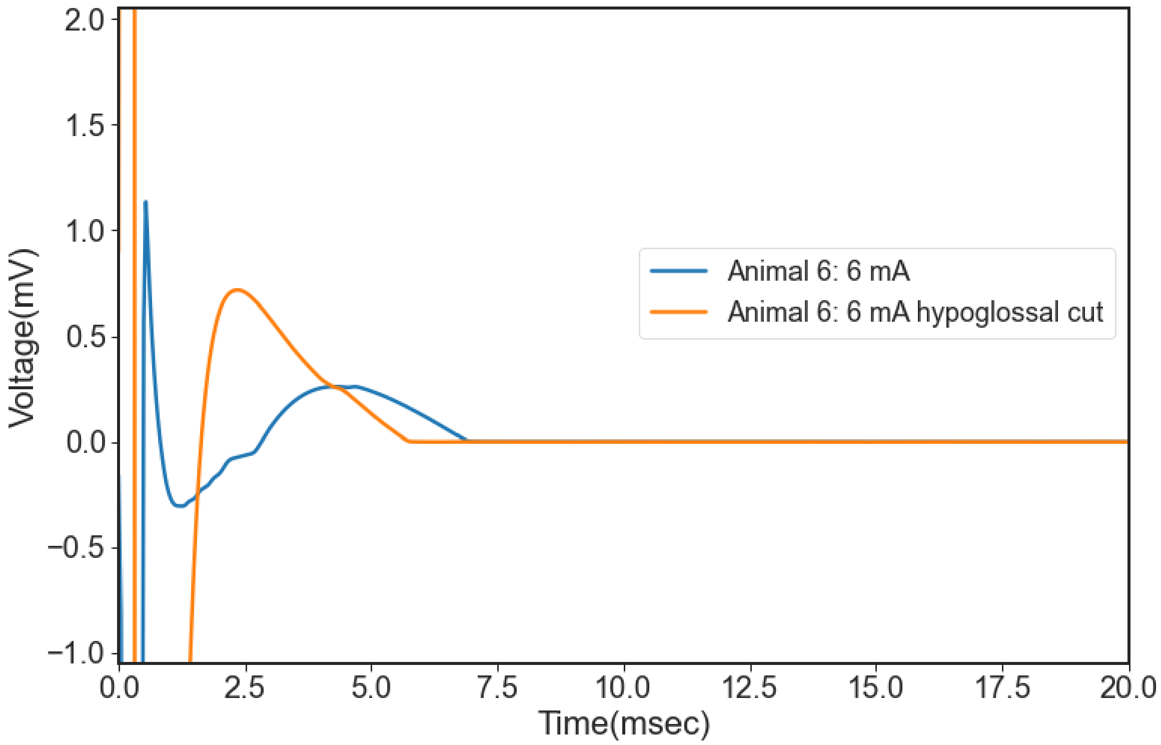


Subject 9

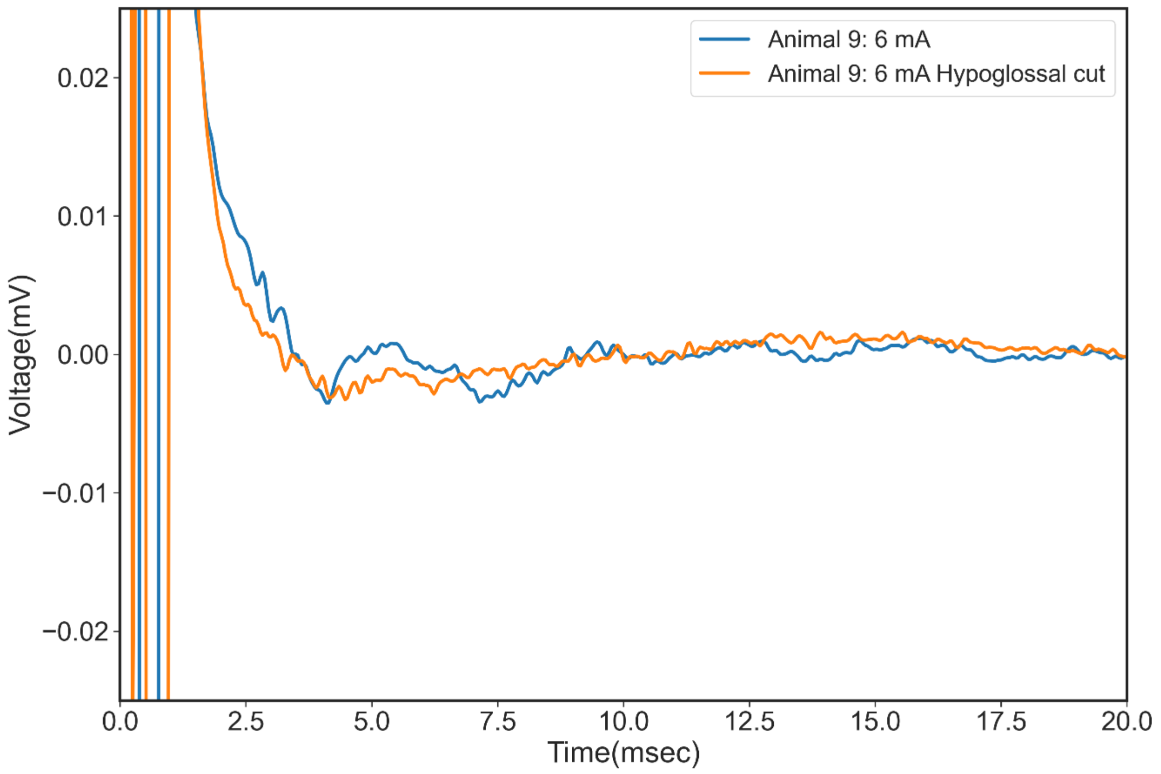


Subject 11

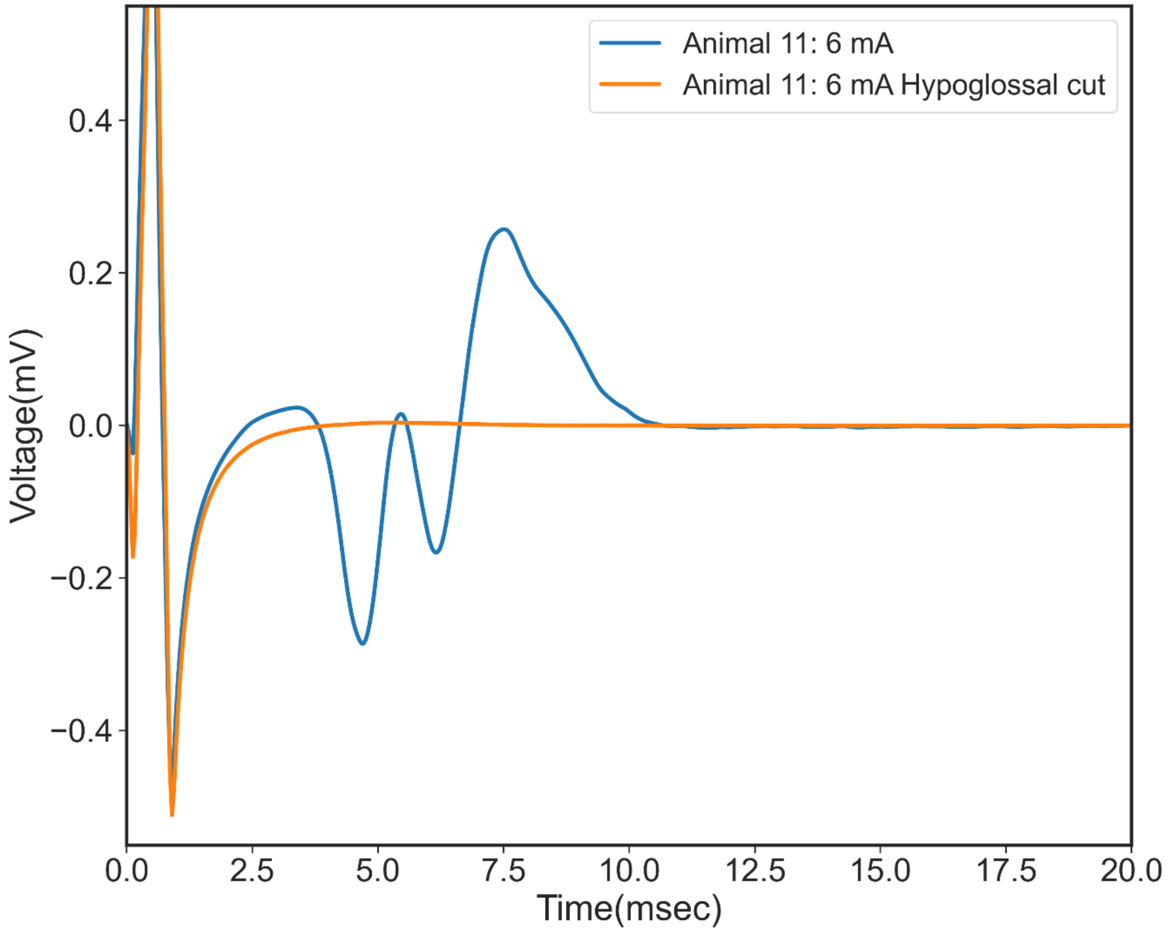


***Glosso-pharyngeal plexus for Constrictors***

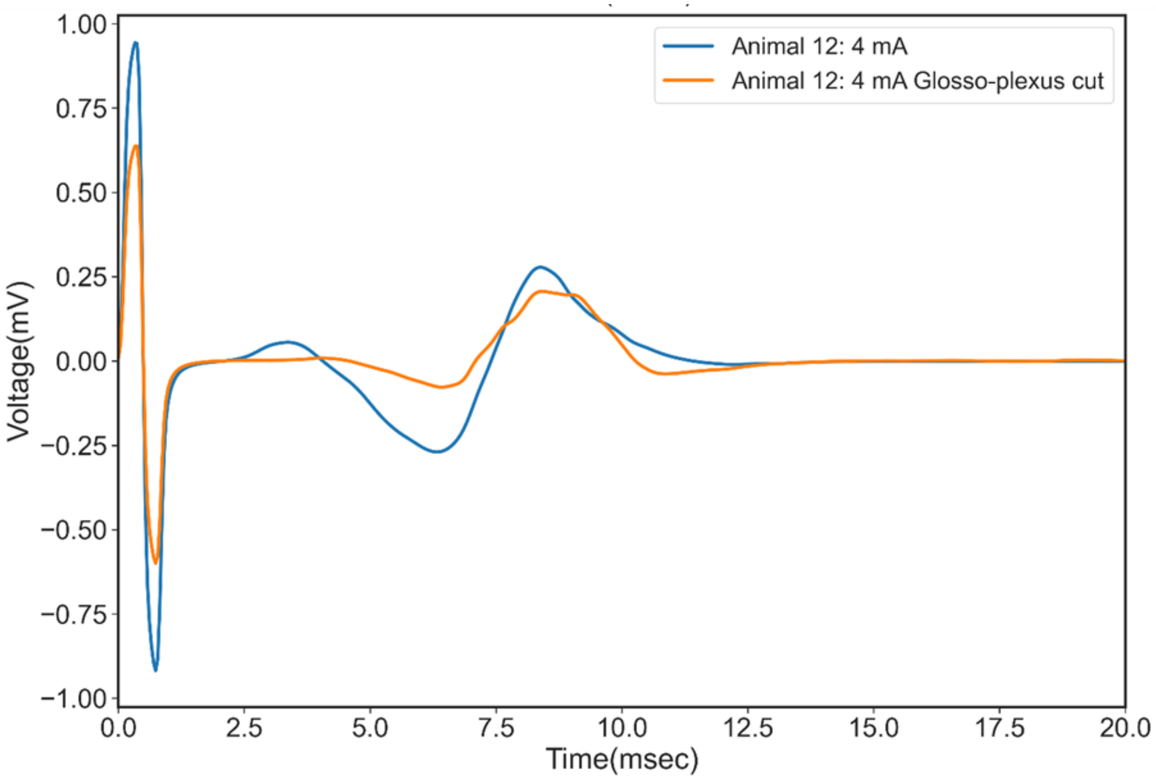


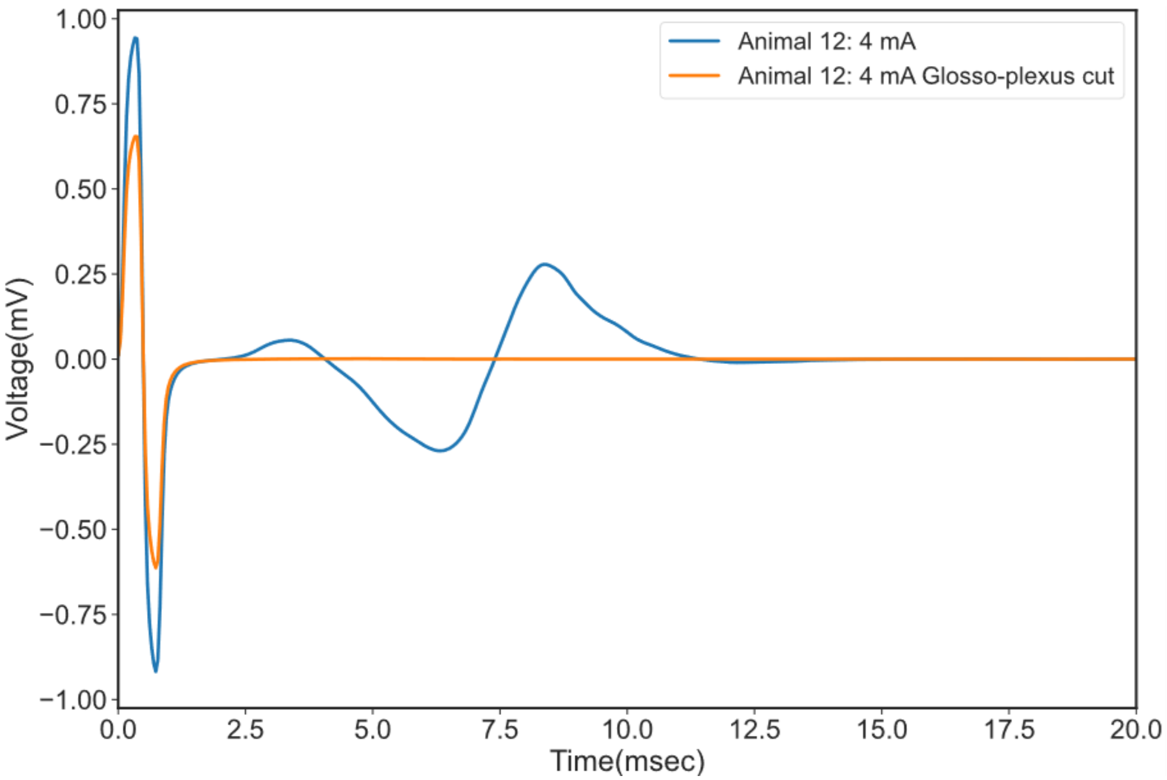


Supplemental Figure 6: Top) EMG signal recorded from the constrictor muscle group post transection of the SLN, RLN and one branch of glossopharyngeal plexus. The recorded EMG was attenuated although not eliminated. Bottom) Constrictor EMG eliminated post transections of the plexus from the glossopharyngeal and the pharyngeal branch of the vagus nerve. These data pointed to one single point of nerve activation but putatively multiple small branches innervating the same muscle.

**Trapezius muscle instrumentation issues**

The trapezius muscle is responsible for shoulder movement and is difficult to instrument and reliability record EMG signals from. The location and constant jostling of the wires due to movement resulted in the EMG wires being pulled out in multiple subjects. As the recording from this target muscle could not be replicated across subjects, it was dropped from the analysis and is not reported as a main off-target muscle although the accessory nerve does innervate it.

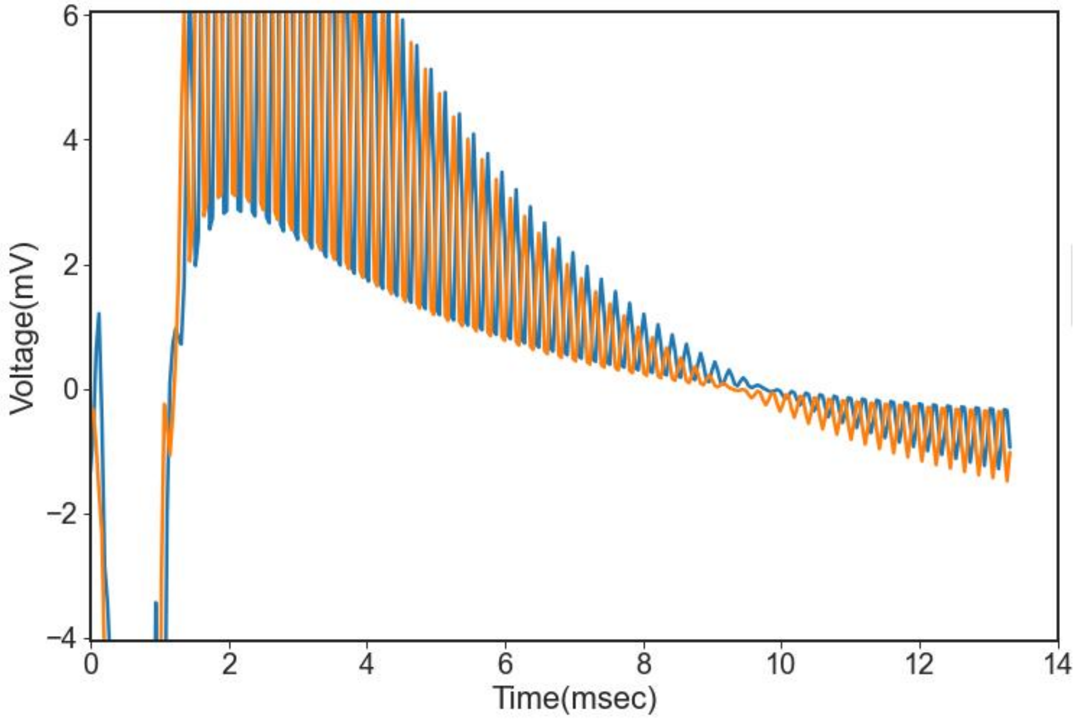


Supplemental Figure 7: Lose wires of the Trapezius EMG resulted in noise and unreliable recordings

**Blood pressure drops**

Subject 10

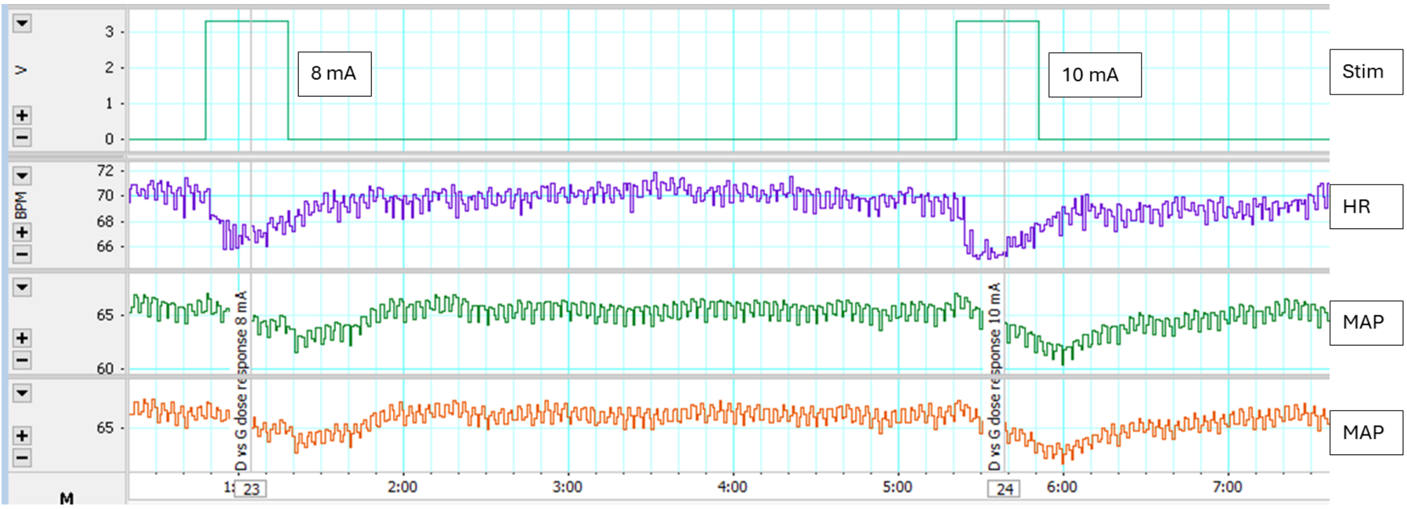


Subject 9

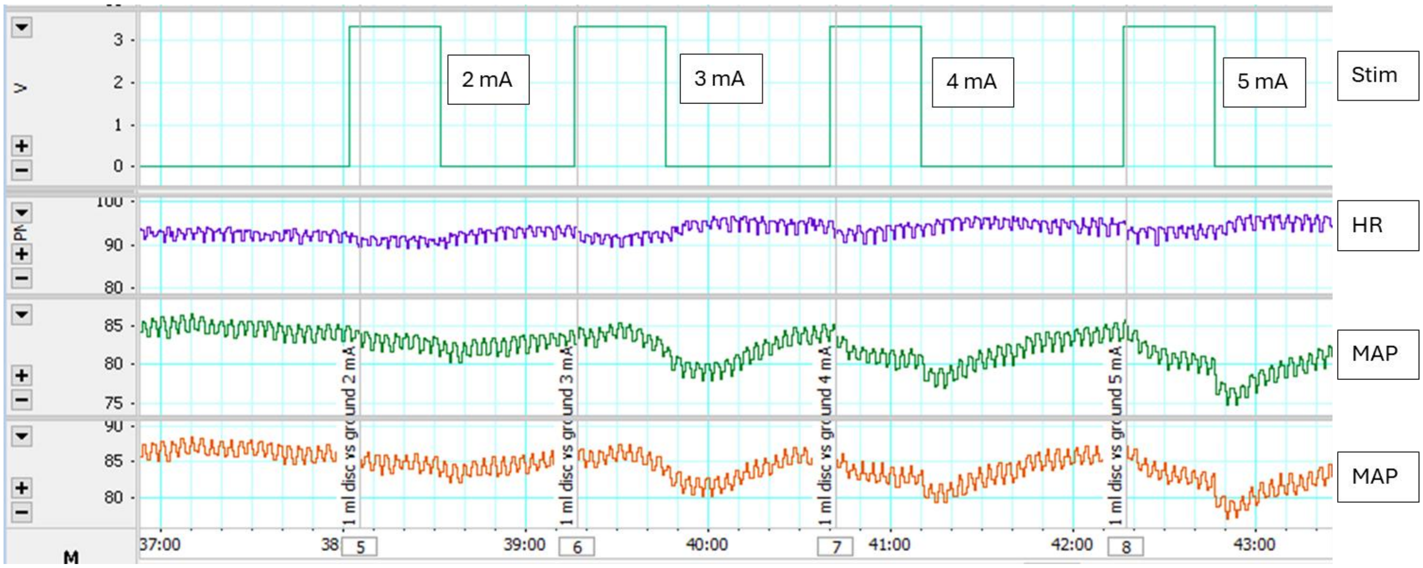


Subject 11

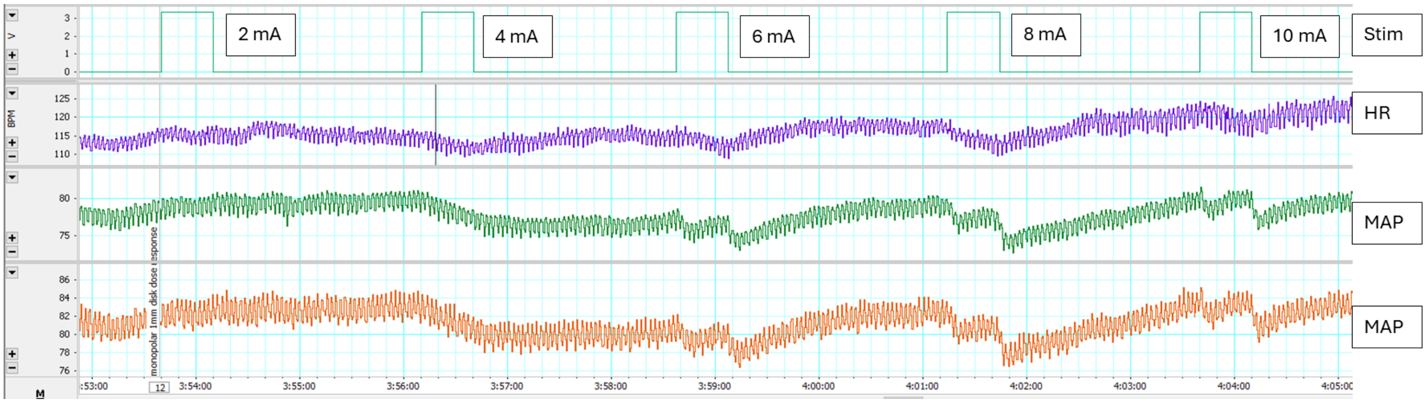


Supplemental Figure 8: Blood pressure drops from additional subjects from cohort collected during dose response curve stimulation.

**Outlier from EMG Dose response curve analysis**

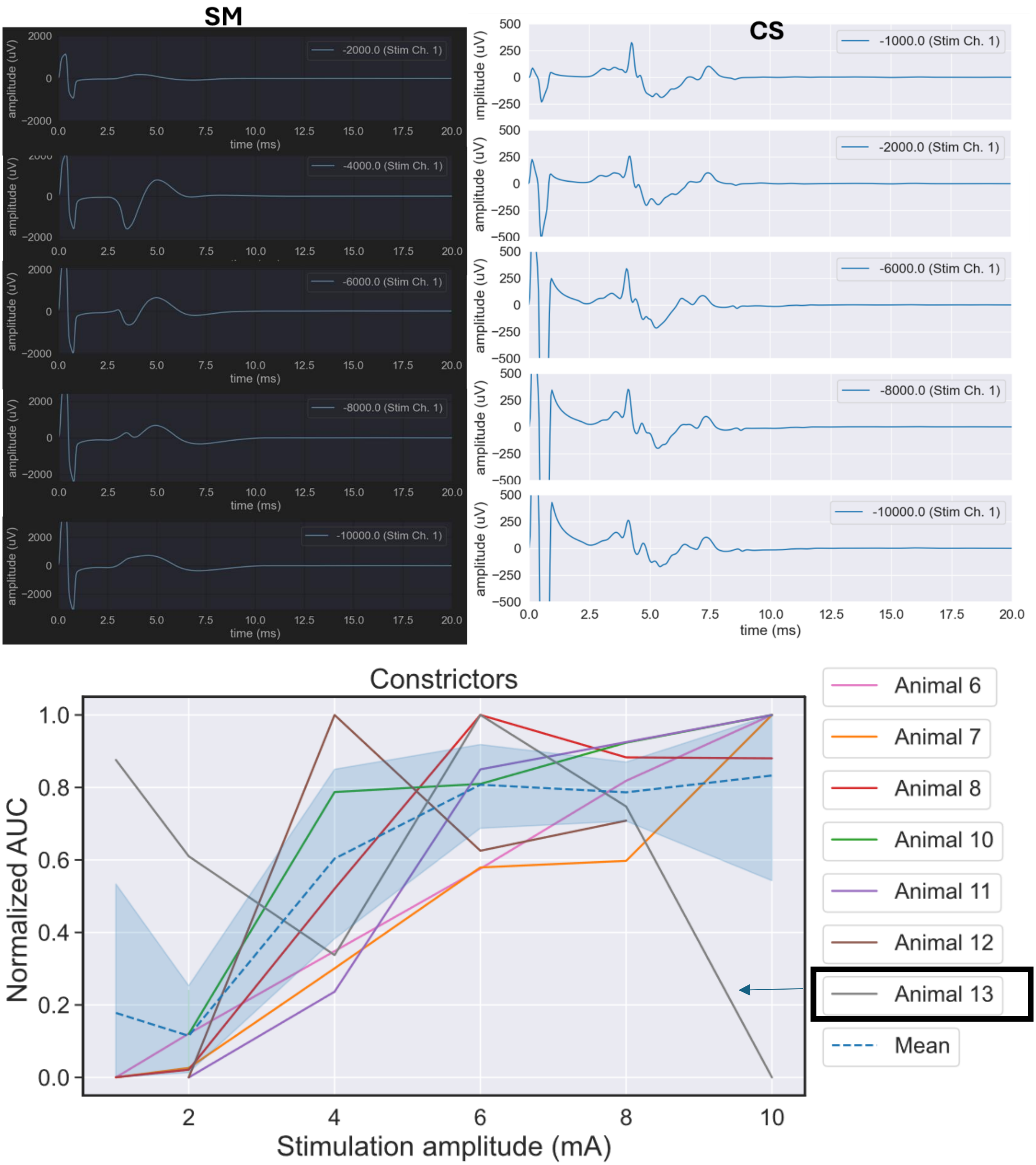


Supplemental Figure 9: Subject 13 had atypical of EMG dose response curve wherein higher amplitudes evoked smaller EMG than lower intensities. There could be multiple reasons for this including instrumentation error in the EMG wires where a large muscle contraction caused the wires to be lose, change in the fluid/edema in the pocket which changed the activation pathway, improper placement of EMG electrodes to record the evoked signal accurately or muscle fatigue.

#### Cohort EMG recordings for selecting activation threshold per muscle

Subject 6

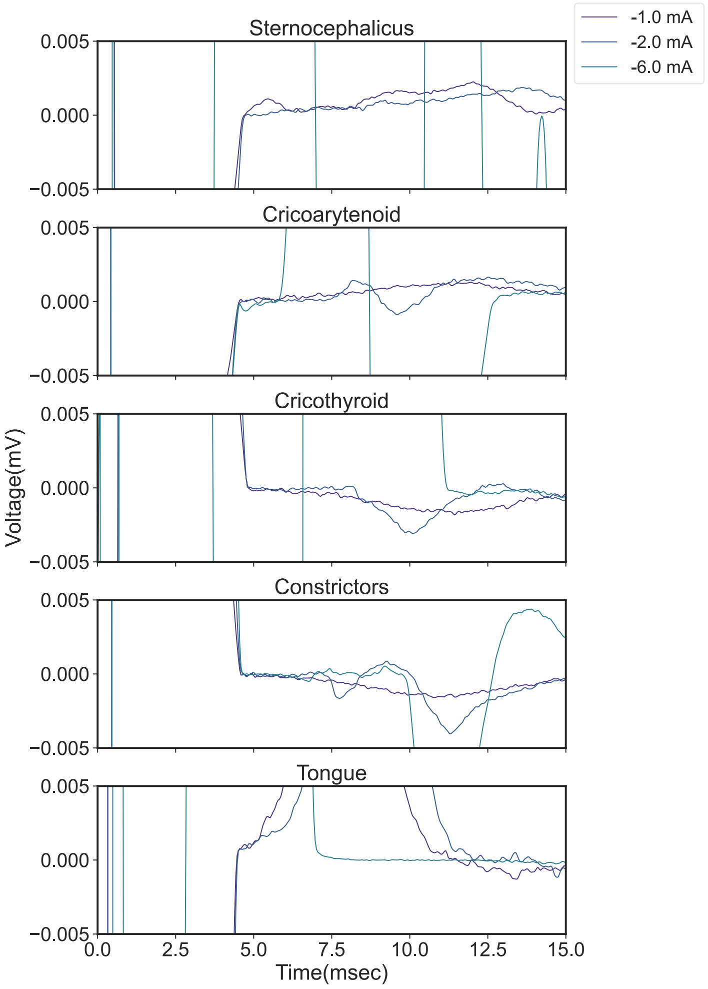


Subject 7

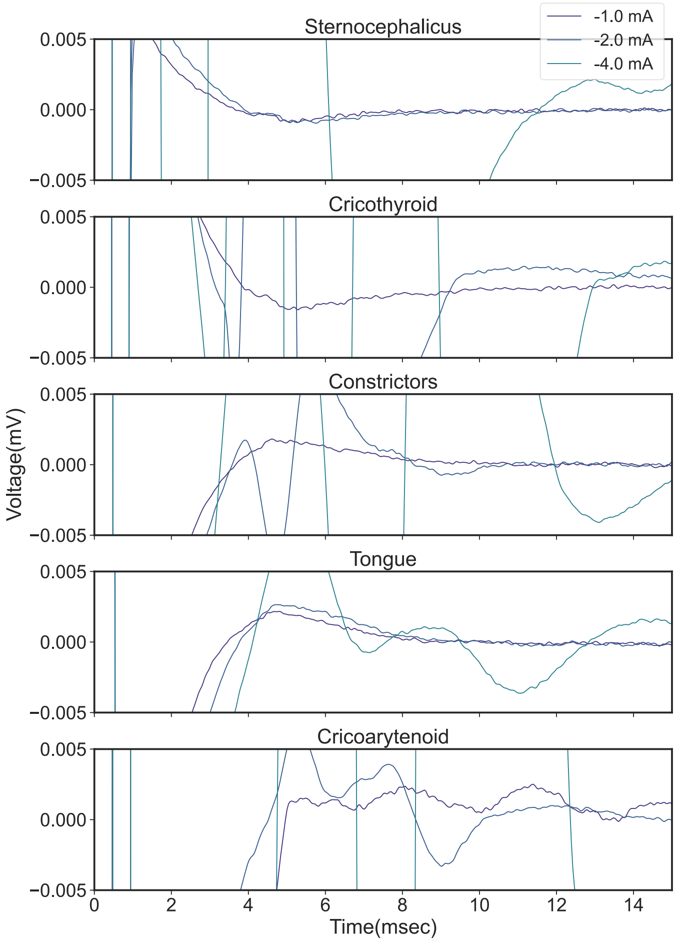

Subject 9

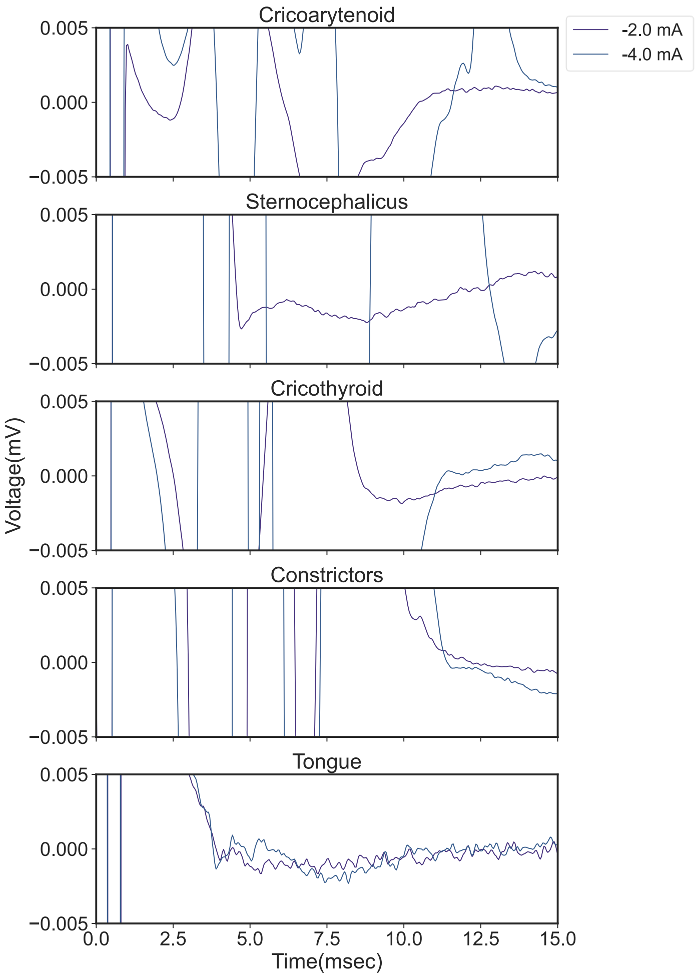


Subject 10

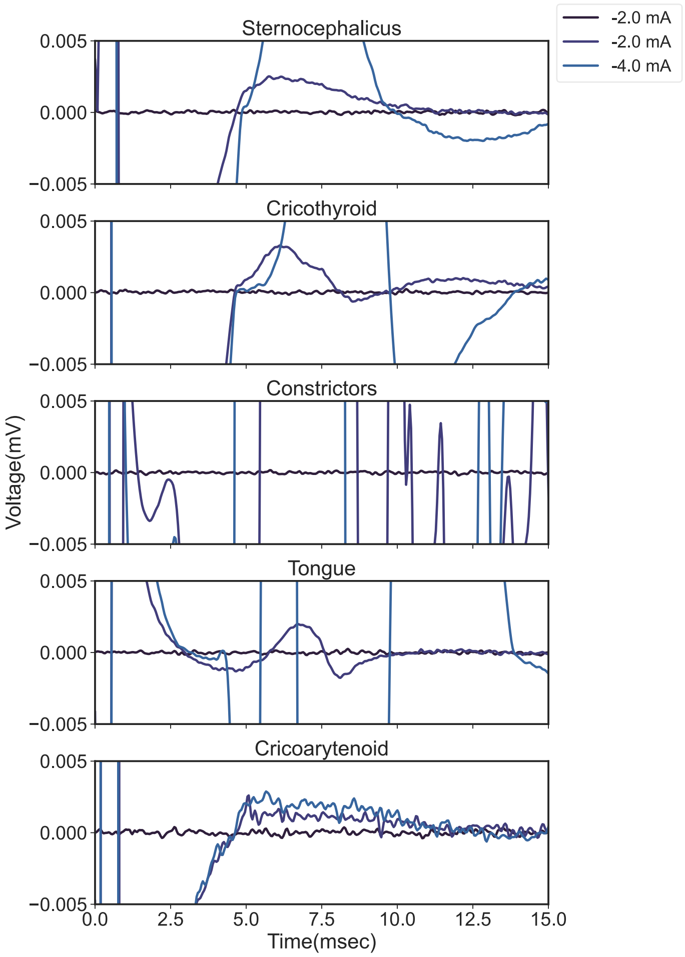

Subject 11

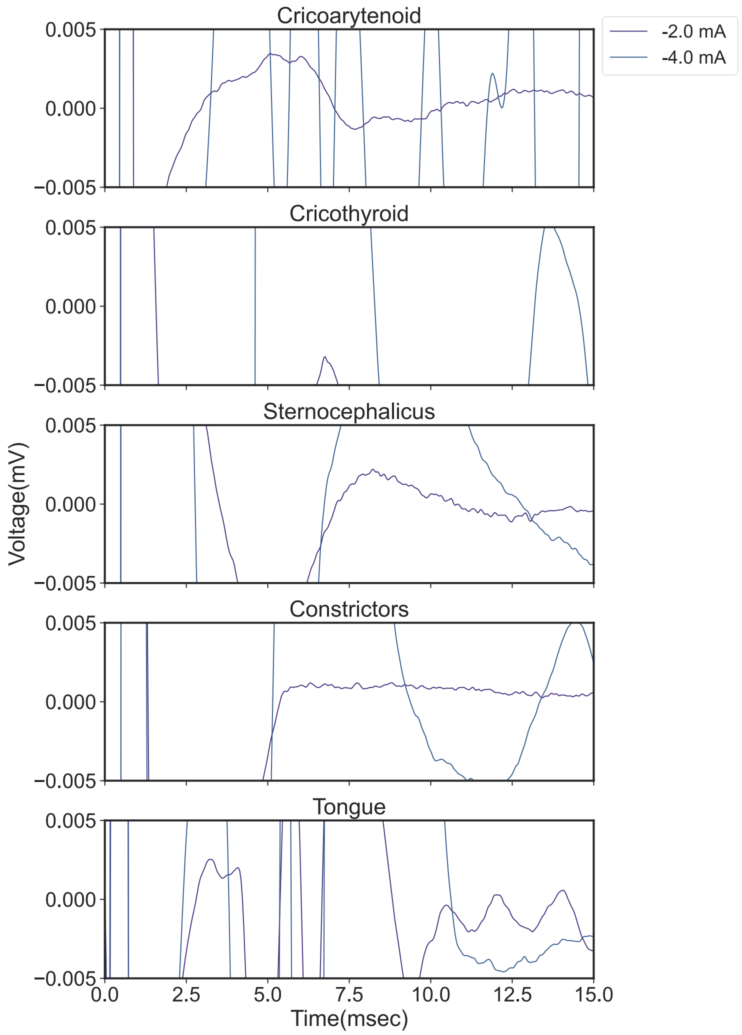

Cricoarytenoid
Cricothyroid
Sternocephalicus
Constrictors
Tongue
-2.0 mA
-4.0 mA
Voltage(mV)
Time(msec)
0.005
0.000
−0.005
0.0
2.5
5.0
7.5
10.0
12.5
15.0


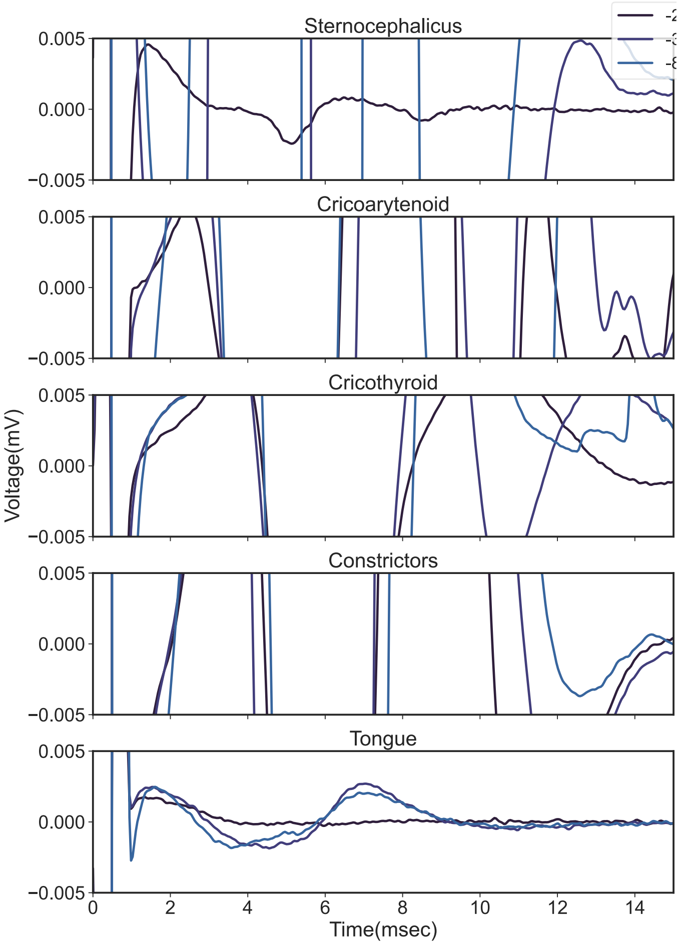

Sternocephalicus
Cricoarytenoid
Cricothyroid
Constrictors
Tongue
Voltage(mV)
Time(msec)
0.005
0.000
−0.005
0
2
4
6
8
10
12
14

Subject 13

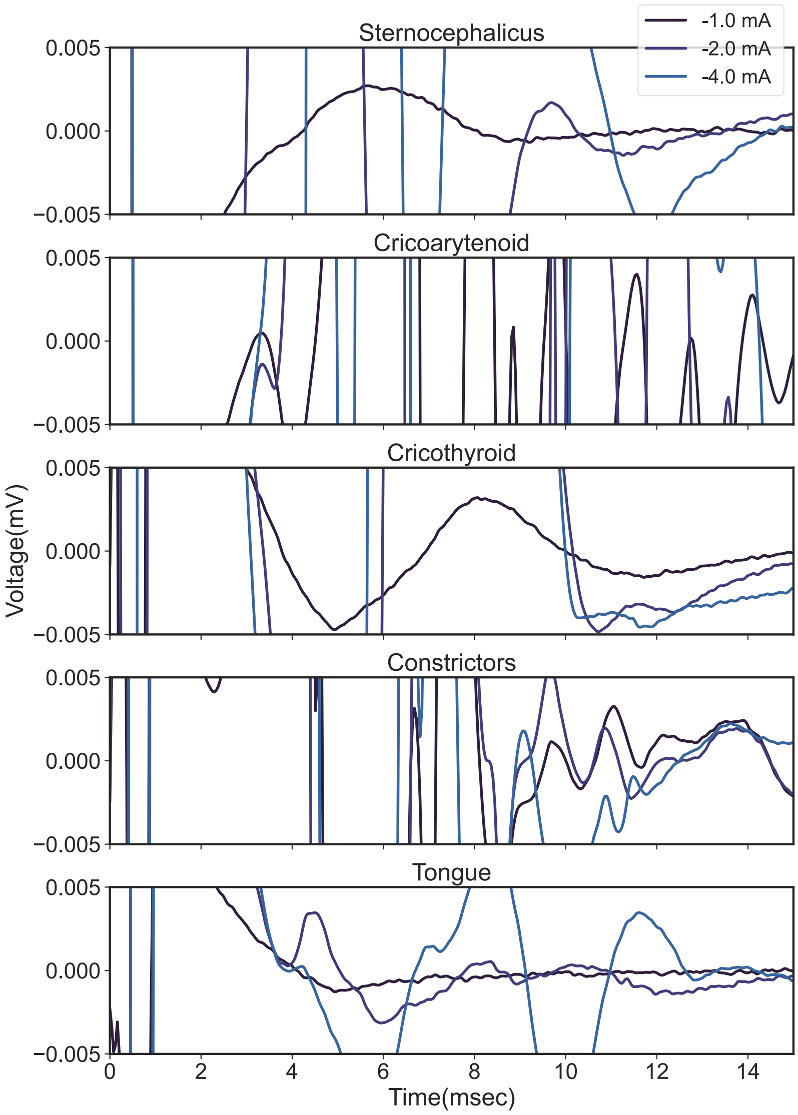


Supplemental Figure 10: Cohort EMG recordings to identify activation thresholds for different muscle groups. If the EMG signal of a particular muscle crossed the amplitude window of 10 uV pk-pk, that stimulation amplitude was noted as the activation threshold for that particular muscle group.
